# Cell Barcoding Reveals Lineage-dependent Outcomes in hiPSC Cardiac Differentiation

**DOI:** 10.64898/2025.12.12.694049

**Authors:** Sogu Sohn, Daylin Morgan, Cody Callahan, Katelyn Dockery, Amy Brock, Janet Zoldan

## Abstract

Human induced pluripotent stem cell-derived cardiomyocytes (hiPSC-CMs) have potential applications in treating cardiovascular disease but are currently limited in their clinical translation. A primary limitation is the poor clinical scalability of hiPSC-CMs, with the heterogeneity of hiPSC cardiac differentiation significantly contributing to this limitation. We hypothesize that clinical scalability can be improved by tracking and controlling hiPSC clonal heterogeneity, a variable often overlooked in current differentiation approaches. “Fate priming”, wherein clonal lineage identity determines differentiation fate, has been demonstrated in other stem cell differentiation pathways. We investigated fate priming in hiPSC cardiac differentiation using the ClonMapper cell barcoding platform to label, track, and isolate distinct hiPSC lineages from the same cell line. We show that certain hiPSC lineages preferentially differentiate into hiPSC-CMs or non-CMs. After isolating lineages with apparent fate priming, we found significant differences in cardiac differentiation outcomes between these single-clone populations and heterogeneous, multi-clone hiPSC populations. These findings indicate that lineage identity influences hiPSC cardiac differentiation outcomes.

**SIGNIFICANCE STATEMENT:** Cardiovascular disease is a significant global health concern that can be addressed by engineering artificial tissues to develop new treatments for heart disease or to directly replace damaged heart tissue. Stem cells are a useful tool for engineering these tissues because of their ability to become cardiomyocytes. However, their clinical translation is limited by variability in the process of differentiating stem cells into cardiomyocytes. This article reports findings that show different lineages of genetically identical human induced pluripotent stem cells have different capacities for differentiating into cardiomyocytes, which may contribute to the variability observed.

## INTRODUCTION

Cardiovascular disease (CVD) remains a leading cause of death and disability, causing 20.5 million deaths and 422 million disability-adjusted life years globally in 2021^1^, making it critical to improve methods for preventing and treating CVD. Human induced pluripotent stem cells (hiPSCs) are a valuable tool in this endeavor, as hiPSCs can be differentiated into cardiomyocytes (hiPSC-CMs) and used to engineer tissues that mimic native human heart physiology^2^. While these engineered tissues have potential to replace damaged tissue, model CVD progression, or evaluate novel interventions for treating CVD^3,4^, hiPSC-CM clinical translation faces limitations.

Although the immaturity of hiPSC-CMs is a well-researched limitation, an overlooked but significant limitation is the heterogeneity of hiPSC cardiac differentiation, which generates non-CMs alongside CMs of varying functional maturity and lacks precise control over CM subtype specification^5,6^. Because cardiac differentiation is time- and labor-intensive, inadequate control over CM yield, functionality, and subtype greatly limits the scalability of hiPSC-CMs. Approaches like fluorescence activated cell sorting (FACS) and glucose starvation isolate hiPSC-CMs to establish pure hiPSC-CM populations^7,8^. However, these approaches still produce off-target cell types that are discarded, resulting in wasted resources. Accordingly, there is a clear need to elucidate mechanisms that regulate hiPSC cardiac differentiation and improve control over the process to reduce heterogeneity and improve clinical relevance of hiPSC-CMs.

We hypothesize that clonal lineage-specific responses to cardiac differentiation contribute to heterogeneous outcomes. Due to self-renewal, hiPSC populations comprise subpopulations of clonal lineages containing cells descended from the same parent cell. Epigenetic differences between lineages may contribute to different lineages being preprogrammed/primed to differentiate into different phenotypes, with lineage-dependent fate priming reported in hematopoietic stem cells^9,10^ and reprogramed mesenchymal stem cells^11^ through use of cellular barcodes. Accordingly, we hypothesize that cell barcoding can be similarly applied to hiPSC cardiac differentiation. Cell barcoding involves inserting unique nucleotide sequences into cell genomes, resulting in cells being labeled with unique, heritable genetic barcodes that can be used to track changes through multiple cell divisions^12^. This means that hiPSCs generated via self-renewal after barcoding will retain the same barcode as their parent cell, allowing sequencing to be used to identify which hiPSCs belong to the same clonal lineage. This barcoding approach enables tracking of hiPSCs from terminally differentiated states, such as hiPSC-CMs, back to their undifferentiated origins. Performing this lineage tracing for an entire barcoded hiPSC population could determine when clonal lineages in the population diverge and differentiate towards CM and non-CM fates.

Herein we report our use of the ClonMapper cell barcoding platform to track hiPSC lineages. A novel feature of ClonMapper is that barcode labels are embedded in a functional single-guide RNA (sgRNA) sequence that facilitates live, sequence-specific recall of barcoded cells for further experimentation and analysis, as opposed to the destructive isolation methods used by other cell barcoding platforms^13^. We established a population of ClonMapper-barcoded hiPSCs that remained pluripotent and able to differentiate into hiPSC-CMs and identified different lineages that were preferentially enriched in CMs and non-CMs. Two lineages enriched in non-CMs were successfully isolated with ClonMapper’s lineage recall, with both isolated populations having worse cardiac differentiation outcomes compared to the parent barcoded population, with sequencing revealing potential key genes to examine when investigating cardiac differentiation. Our results indicate that different hiPSC clonal lineages originating from the same cell line have varying cardiac differentiation potential and that ClonMapper can be used to identify and isolate these lineages, with isolated populations retaining their differentiation potential. This highlights the value of examining hiPSC clonal heterogeneity in cardiac differentiation and the potential for functional cell barcoding methods like ClonMapper to be used to monitor and control clonal heterogeneity to potentially improve cardiac differentiation outcomes.

## RESULTS

### hiPSCs Can be Labeled with ClonMapper Barcodes

hiPSCs were transduced with ClonMapper barcodes following a viral titer assay to determine viral load required to maintain a multiplicity of infection (MOI) ≤ 0.1 (**Fig. S1**), as an MOI ≤ 0.1 results in <0.47% probability of cells being transduced with more than one barcode. This MOI was necessary to prevent single clonal lineages from appearing as multiple different clonal lineages in sequencing analyses. hiPSCs were successfully barcoded with an MOI ≤ 0.1 and sorted using blue fluorescent protein (BFP) signal to generate ClonMapper-barcoded hiPSC populations (**Fig. 1A**), as ClonMapper barcodes contain a BFP gene such that barcoded cells fluoresce. Sorted hiPSCs maintained BFP over several passages (**Fig. 1B-D**), indicating that ClonMapper barcodes can be integrated and retained in hiPSCs.

**Figure 1:**
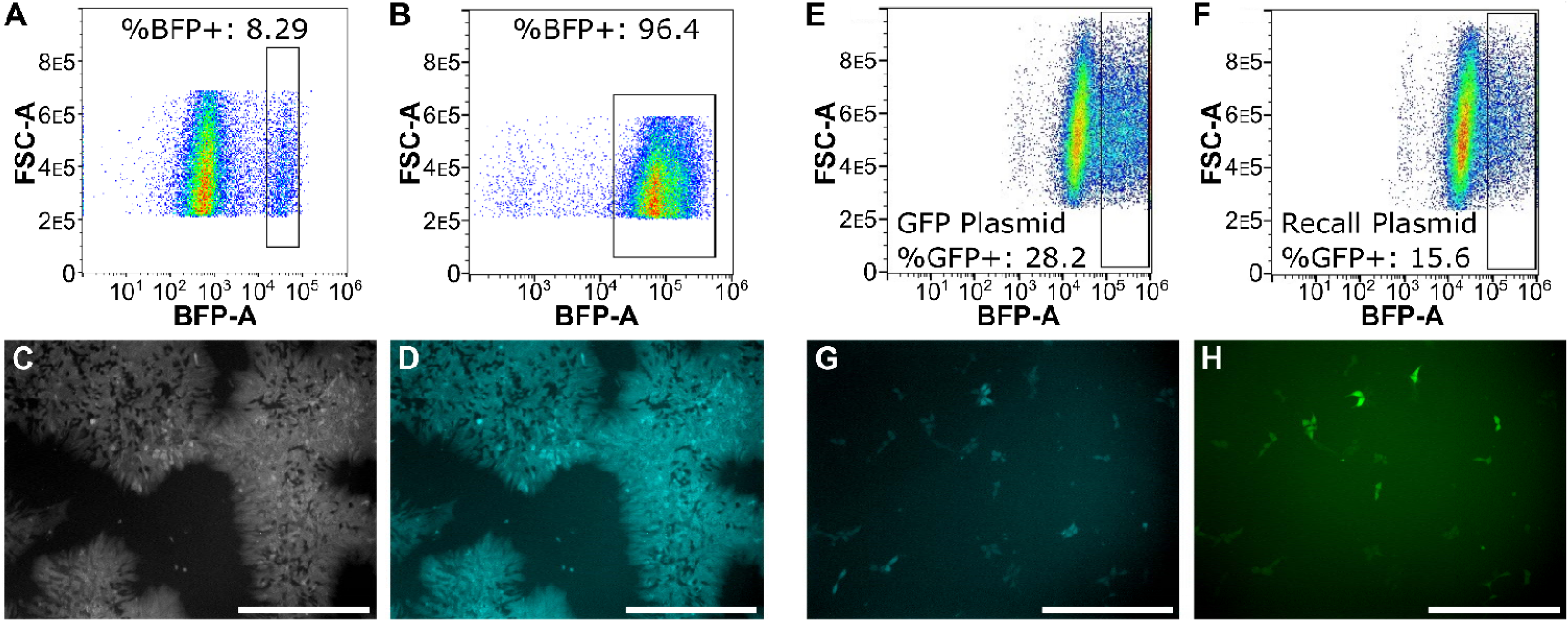
Successful adaptation of ClonMapper barcoding to hiPSCs. hiPSCs were sorted with %BFP+ < 10%, indicating transduction of the barcoded population was successfully performed with an MOI < 0.1 (**A**). ClonMapper barcoded, sorted hiPSCs after being continually expanded and passaged, retaining BFP signal and purity (**B**). Representative images demonstrating that sorted barcoded hiPSC express BFP at levels detectable via fluorescence microscopy, with phase contrast imaging of hiPSC colonies (**C**) overlapping with BFP channel imaging (**D**). Comparing transfection efficiency of ClonMapper barcoded RC hiPSCs sorted after transfection with a GFP plasmid (**E**) and dual recall plasmid and dCas9-VPR plasmids (**F**) reveals hiPSC recall transfection does not require additional optimization for efficient transfection. Recalled hiPSCs survived sorting and adhered to plates, with BFP signal (**G**) and GFP signal (**H**) overlap showing that successfully sorted hiPSCs expressed GFP due to interactions between ClonMapper barcode, recall plasmid, and dCas9-VPR plasmid. All scale bars are 400 µm.

### Barcoded hiPSCs can be Recalled

ClonMapper can uniquely non-destructively isolate single lineages from a barcoded population via its “recall” function (**Fig. S2**). To evaluate ClonMapper recall in hiPSCs, a “recall control” (RC) hiPSC population was generated, wherein all hiPSCs were transduced with identical barcode sequences so all barcoded and sorted hiPSCs contained the same barcode. Sorted RC hiPSCs were then transfected either with two plasmids needed for ClonMapper recall (dCas9-VPR and recall plasmids) or with a single GFP plasmid (to control for baseline transfection efficiency in barcoded hiPSCs), while maintaining equal amounts of total plasmid DNA. Flow cytometry revealed RC hiPSCs that underwent recall transfection had approximately half the %GFP+ signal of RC hiPSCs transfected with the GFP plasmid (**Fig. 1E, F**). Because GFP fluorescence actuated by ClonMapper recall requires simultaneous transfection with two plasmids in contrast to GFP fluorescence actuated by transfection with one GFP plasmid (**Fig. S2**), this demonstrated that ClonMapper recall was half as efficient as GFP plasmid transfection, suggesting no barriers to ClonMapper’s recall in barcoded hiPSCs. Recalled hiPSCs successfully adhered to vitronectin-coated plates, and expressed overlapping GFP/BFP signal, confirming that sorted hiPSCs were isolated based on recall-actuated GFP expression (**Fig. 1G, H**). Transient GFP expression disappeared within 48-72 hours while BFP expression was retained. Taken together, these results demonstrate that ClonMapper’s unique method of non-destructive isolation of single lineages can be performed on hiPSCs.

### Barcoded hiPSCs are Pluripotent and Differentiate into CMs

Expression of pluripotent markers Oct4 and SOX2 (**Fig. S3A,B**) and the ability to differentiate into cTnT-expressing hiPSC-CMs (**Fig. S3C**) demonstrated that barcode hiPSCs remained pluripotent. Furthermore, expression of a GCaMP6f transgene was retained in hiPSCs after barcoding such that hiPSC-CM contractions coincided with fluorescence^14^, allowing fluorescent signal to be used to quantify excitation-contraction coupling (ECC) (**Fig. S3D-F**), as shown in our previous work^15^, demonstrating that ClonMapper did not interfere with existing transgene activity. Barcoded and non-barcoded hiPSC-CMs were co-stained with cardiac markers SIRPα^16^ and cTnT to compare hiPSC-CM yield, with flow cytometry showing no significant difference in %cTnT+/SIRPα+ cells between barcoded and control samples (**Fig. 2A**), indicating barcoding did not lower hiPSC-CM yield. Next, quantifying ECC metrics showed no significant difference between barcoded and non-barcoded hiPSC-CMs in the full-width half-max (FWHM), median-absolute-deviation of time-of-peak-arrival (MAD-TPA), or decay constant of fluorescent signal based on analysis with our MATLAB image analysis pipeline (**Fig. 2B, S4**), indicating barcoding did not worsen hiPSC-CM ECC. Lastly, we performed qPCR to examine differences in relative expression of cardiac genes between barcoded and non-barcoded hiPSCs and hiPSC-CMs, with foldchange expression normalized to the non-barcoded sample at each timepoint (i.e., fold-change values for control are all ∼1). We found no significant difference in expression of myofibrillar gene troponin I 1 (*TNNI1*) in hiPSCs or hiPSC-CMs, and while day 0 barcoded hiPSCs exhibited significantly higher *TNNI3* expression (2.64-fold), there were no significant differences in *TNNI3* expression in early (day 7) or late (day 14) hiPSC-CMs. Next, we examined genes related to calcium storage and action potential duration: ryanodine receptor 2 (*RYR2*), calcium voltage-gated channel subunit alpha1 C (*CACNA1C*), and calsequestrin 2 (*CASQ2*). *RYR2* expression was significantly lower in day 0 barcoded hiPSCs (0.64-fold), however day 7 barcoded hiPSC-CMs had significantly higher *RYR2* expression (1.43-fold), while there was no difference on day 14, suggesting that differences in *RYR2* expression were transient. *CASQ2* and *CACNA1C* expression was not significantly different on day 0, but barcoded hiPSC-CMs had significantly higher expression of both genes on day 7 (*CASQ2*: 1.65-fold; *CACNA1C*: 2.04-fold) and day 14 (*CASQ2*: 2.95-fold; *CACNA1C*: 1.91-fold). *CACNA1C*, and *CASQ2* are involved in handling and storing calcium ions^17,18^, with greater expression of these genes being associated with greater action potential duration and calcium storage capacity. Collectively, these data indicate that ClonMapper barcoding did not reduce CM yield or functionality. Although several calcium handling genes showed differential expression between barcoded and unlabeled hiPSC-CMs, given the lack of functional differences in ECC analysis, control and barcoded hiPSC-CMs can be treated as equivalent.

**Figure 2:**
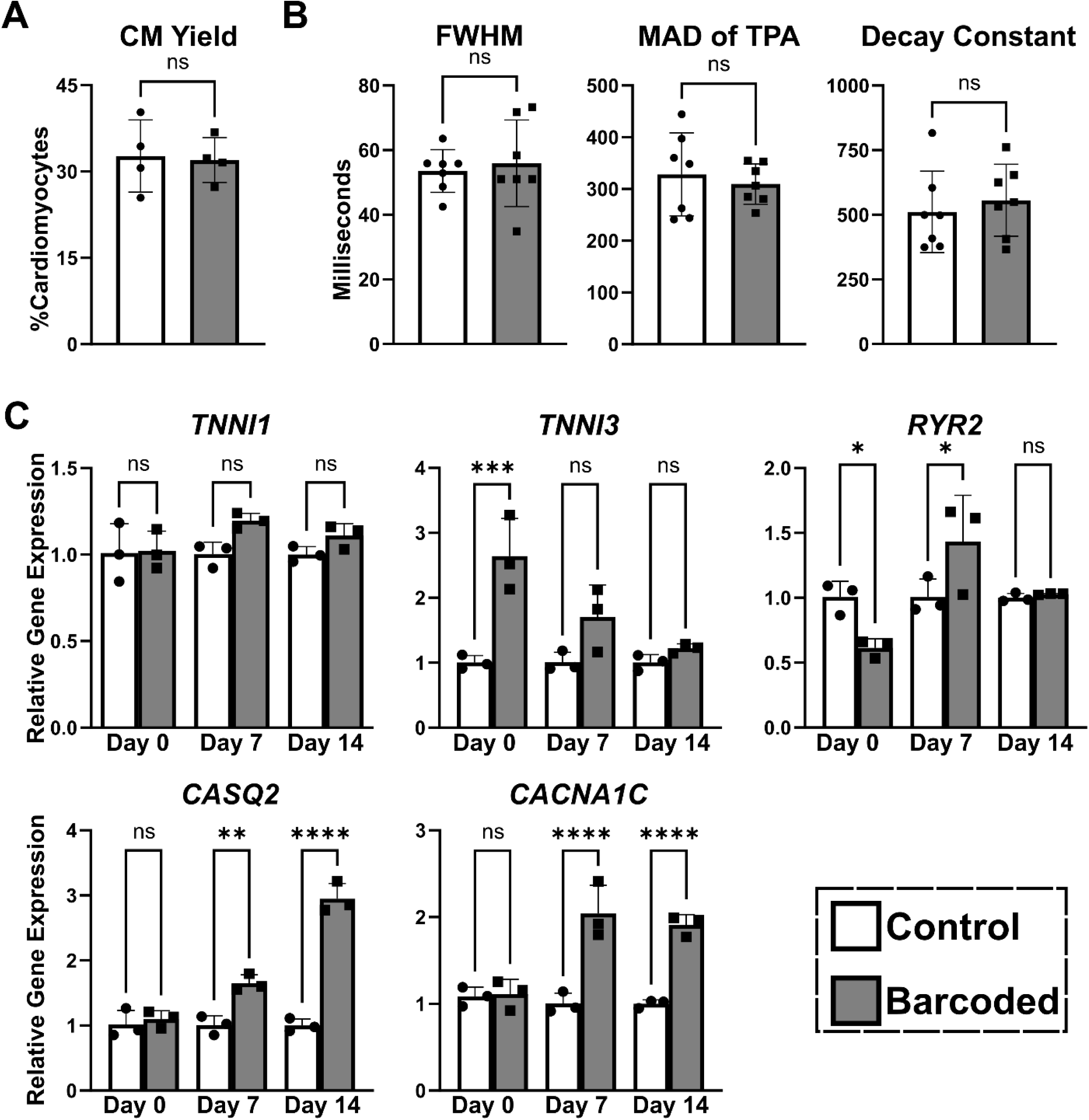
Comparisons of cardiac differentiation outcomes of non-barcoded and barcoded hiPSCs. Flow cytometry data of hiPSCs stained for surface marker SIRPα or intracellular cTnT revealed no significant difference in relative abundance of hiPSC-CMs after differentiation (**A**; n = 4, >30,000 events per n). MATLAB imaging pipeline analysis output shows no difference in functional beating metrics for full-width half-max (FWHM), beats per minute (BPM), median absolute deviation (MAD-TPA), and decay constant (Tau) (**B**; n = 6). Real-time qPCR data quantified using the ΔΔC_T_ approach revealed no significant differences that arise in relative expression of myofibrillar genes (*TNNI1*, *TNNI3*) and some significant increases in expression of calcium-handling genes (*RYR2*, *GJA1*, *CACNA1C*, *CASQ2*) in barcoded cells after differentiation into hiPSC-CMs (**C**; n=3). Data are represented as mean ± SD; ** indicates *p* ≤ 0.01, *** indicates *p* ≤ 0.001.

### Cardiac Differentiation Significantly Impacts Clonal Dynamics

Flow cytometry revealed a significant reduction in BFP+ cells after day 3 of cardiac differentiation (**Fig. S5A**), with similar reductions in non-GCaMP6f WTC11 hiPSC cardiac differentiation (**Fig. S5B**) and DF19-9-11T.H hiPSCs (WiCell) endothelial progenitor differentiation^19^ (**Fig. S5C**). To determine whether this was the result of barcode silencing or removal, gDNA was extracted from BFP+ and BFP- barcoded hiPSC-CMs sorted by FACS, amplified via PCR reaction targeting ClonMapper barcode sequences, then electrophoresed in agarose gel (**Fig. S6)**. This showed ClonMapper barcodes were still present in BFP- cells, enabling barcode quantitation using high-throughput sequencing.

Next, we established a parental barcoded (PB) hiPSC population wherein baseline lineage abundance was quantified via next-generation sequencing, providing a defined starting point essential for accurately interpreting lineage dynamics. The selected PB population had a high barcode heterogeneity (Shannon Diversity index >5) and an even barcode distribution (most abundant barcodes comprised ∼2% of the population) (**Fig. S7A**). This indicated that the PB population was viable for capturing clonal dynamics through differentiation, as resolving changes in clonal dynamics would be difficult in a population with a skewed barcode distribution and low diversity. Abundance sequencing and analysis performed on gDNA extracted from differentiating PB cells throughout differentiation revealed that the PB population undergoes statistically significant reductions in Shannon diversity (**Fig. S7B**) and notable changes in abundance of many lineages through differentiation (**Fig. S7C**).These findings demonstrate that ClonMapper can be used to track changes in hiPSC lineage abundance through cardiac differentiation, with some of these changes being significant.

### Specific Lineages are Enriched in Cardiomyocytes and Non-myocytes

We examined the relationship between lineage identity and differentiation trajectory by sorting PB cells positive or negative for CD140a (cardiac mesoderm marker^20^) on day 5 and SIRPα (hiPSC-CM marker^16^) on day 14 to isolate subpopulations that succeeded or failed to become cardiac mesoderm cells and hiPSC-CMs (**Fig. S8A**). Abundance sequencing of gDNA from each subpopulation (CD140a+, CD140a-, SIRPα+, SIRPα-) was used to quantify changes in lineages’ log2(counts per million) (log2(CPM)) in each subpopulation (**Fig. S8B**). This revealed that lineage abundance varied in each subpopulation, suggesting that lineage abundance was influenced by differentiation trajectory. We then calculated foldchange (FC) of each lineage’s CPM abundance in the sorted populations relative to day 0. Plotting each lineage’s SIRPα+ log2(FC) against its SIRPα-log2(FC) revealed lineage-specific responses to cardiac differentiation (**Fig. 3**). Lineages in quadrant I (log2(FC)_SIRPα+_>0, log2(FC)_SIRPα-_<0) showed increased abundance in CMs and decreased abundance in non-CMs, indicating preferential differentiation into hiPSC-CMs. Lineages in quadrant III (log2(FC)_SIRPα+_<0, log2(FC)_SIRPα-_>0) displayed preferential differentiation into non-CMs. Lineages in quadrant II (log2(FC)_SIRPα+_>0, log2(FC)_SIRPα-_>0) non-preferentially increased in abundance in both CMs and non-CMs, while lineages in quadrant IV (log2(FC)_SIRPα+_<0, log2(FC)_SIRPα-_<0) non-preferentially decreased in abundance. Accordingly, we identified PB hiPSC lineages preferentially enriched in either hiPSC-CMs (quadrant I) or non-CMs (quadrant III), suggesting that hiPSC lineage identity influences their differentiation outcomes. We similarly plotted log2(FC) on day 5 to visualize lineage enrichment in day 5 subpopulations (**Fig. S9**). When we identified lineages enriched in CD140a+ or CD140a- populations on day 5 (indicative of following or diverging from a cardiac trajectory, respectively) and identified the positions of these same lineages on the day 14 log2(FC) plot, many lineages changed quadrants, suggesting that CD140a alone is insufficient to reliably identify differentiating cells committed to a cardiac fate.

**Figure 3:**
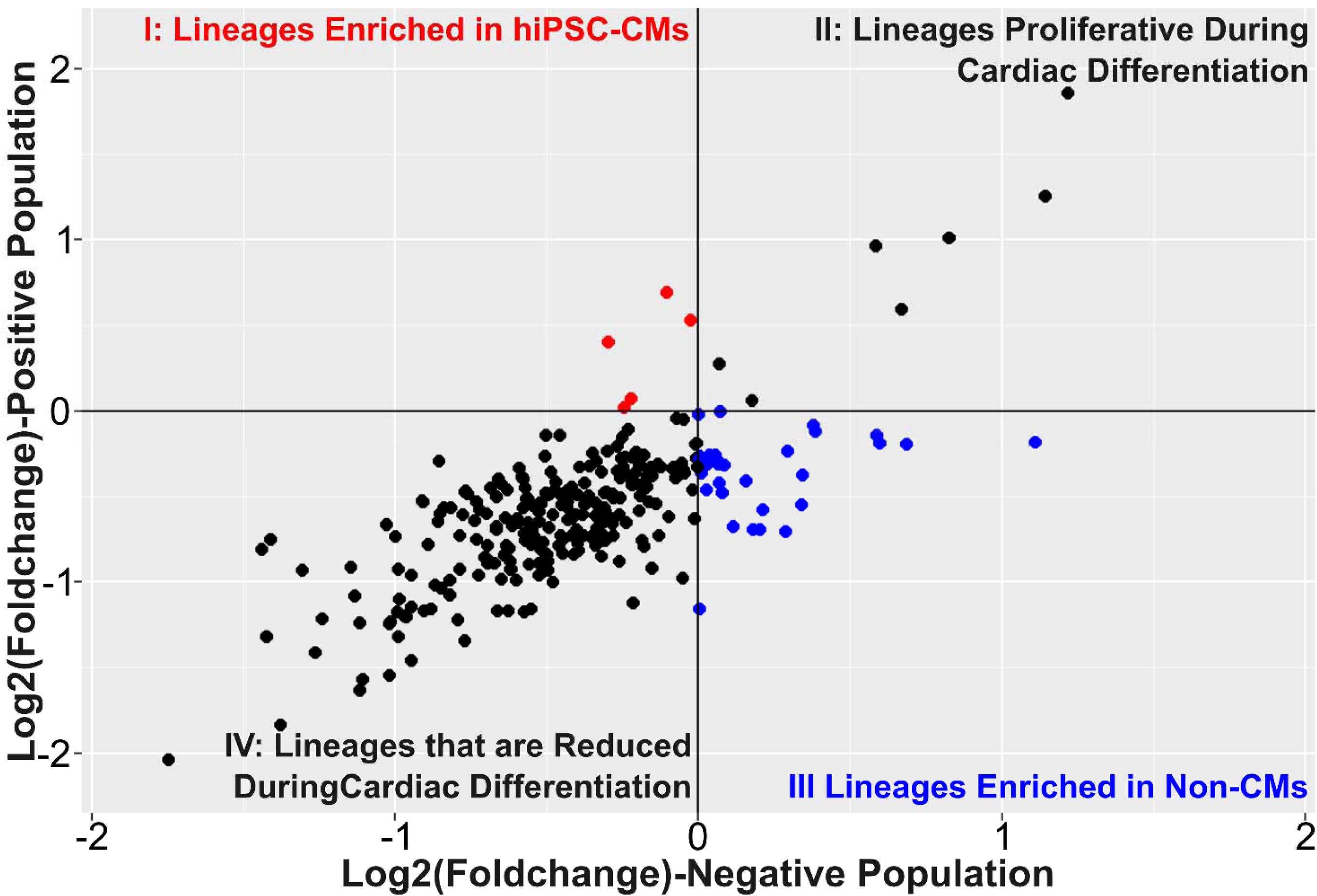
Plot of day 14 Log2(Foldchange) in SIRPα+ vs SIRPα- sorted populations for each PB population lineage. Change in relative abundance of barcodes as determined by average log2(foldchange) in the sorted SIRPα+ and SIRPα- populations on day 14 of differentiation relative to day 0 undifferentiated populations plotted against each other (n = 6). Cardiogenic lineages (red dots, quadrant I) were identified based on positive log2(FC) in SIRPα+ populations and negative log2(FC) in SIRPα- populations, i.e., enrichment i a CM phenotype. Non-cardiogenic lineages (blue dots, quadrant III) were identified based on inverse log2(FC) values, i.e., enrichment in a non-CM phenotype. Quadrants II and IV contain lineages that proliferated well and poorly during cardiac differentiation, respectively.

### ClonMapper Recall can Establish Single-Lineage hiPSC Populations

We used ClonMapper’s recall to isolate PB lineages enriched in hiPSC-CMs and non-CMs to further evaluate their cardiac differentiation potential. We recalled the three most abundant lineages enriched in CMs and the two most abundant lineages enriched in non-CMs. Sanger sequencing of gDNA isolated from the recalled populations revealed that only the two non-cardiogenic lineages were successfully recalled, establishing single-lineage hiPSC populations (**Fig. S10**), with these lineages designated low-differentiation-efficiency 1 (LD1) and LD2. LD1 and LD2 were cultured with no signs of stress or autodifferentiation, indicating that clonally homogeneous hiPSC populations can survive and proliferate after recall. Failure to establish pure, single-lineage populations after recalling the other lineages indicates that there are challenges to recalling hiPSC lineages not predicted by earlier experiments recalling RC hiPSCs.

### Clonal Identity Influences Differentiation Outcomes

Difference in hiPSC-CM yield was quantified by percentage of cTnT+/SIRPα+ cells using flow cytometry. While there was no significant difference in yield between PB (31.99%) and LD1 (36.36%), LD2 (5.78%) yielded significantly fewer hiPSC-CMs (**Fig. 4A**). Thus, while both LD1 and LD2 lineages were enriched in non-CMs, only LD2 hiPSCs exhibited significantly lower hiPSC-CM yield relative to PB when differentiated in isolation.

**Figure 4:**
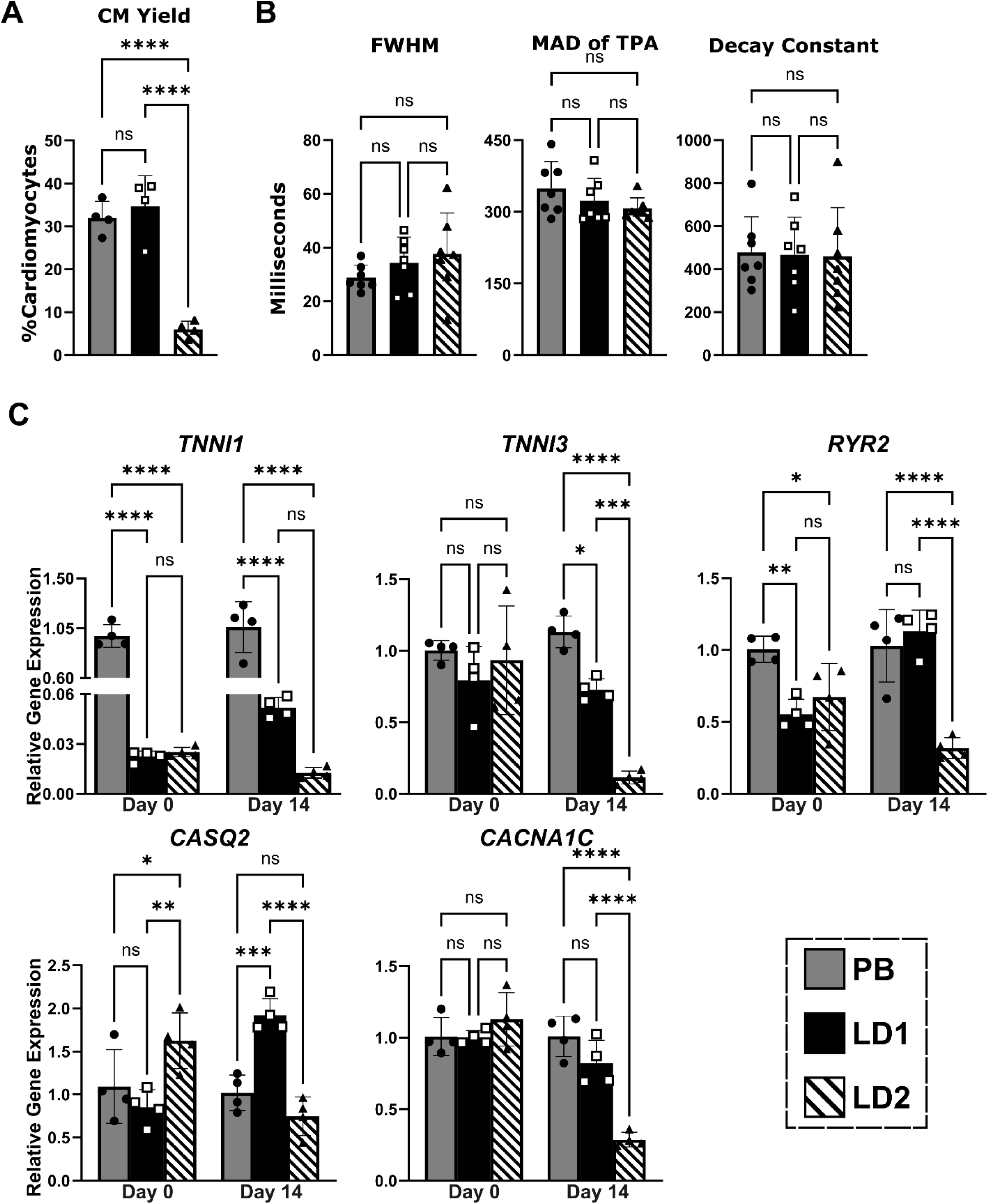
Comparison of cardiac differentiation outcomes between PB, LD1, and LD2 hiPSCs. Flow cytometry data of hiPSCs stained for surface SIRPα or intracellular cTnT (**A**; n = 4, >30,000 events per n). MATLAB imaging pipeline analysis output of excitation-contraction coupling metrics of full-width half-max (FWHM), median absolute deviation (MAD-TPA), and decay constant (Tau) (**B**; n = 4). Real-time qPCR data quantified using the ΔΔC_T_ approach, comparing relative gene expression in day 0 undifferentiated and day 14 differentiated populations with foldchange determined using PB as the reference sample for both timepoints. Data are represented as mean ± SD; * indicates *p* ≤ 0.05, ** indicates *p* ≤ 0.01, *** indicates *p* ≤ 0.001, **** indicates *p* ≤ 0.0001.

Comparing ECC of PB, LD1, and LD2 hiPSC-CMs, we observed no significant differences in FWHM, MAD-TPA, or decay constant between all three populations (**Fig. 4B**). However, while not statistically significant, LD1 and LD2 both had lower average FWHM and higher average MAD-TPA values compared to PB, with lower FWHM and higher MAD-TPA values associated with less functionally mature CMs^21^. Thus, while not statistically significant there was some evidence of LD1 and LD2 hiPSC-CMs having poorer functionality relative to PB.

Lastly, we performed qPCR analysis on days 0 and 14 of cardiac differentiation to quantify relative expression of cardiac-specific genes in undifferentiated hiPSCs and late hiPSC-CMs (**Fig. 4C**). qPCR results were analyzed with foldchange expression normalized to PB at each timepoint (i.e., foldchange values are all ∼1 for PB).

#### Day 0

No significant difference was observed between the three populations in relative expression of *TNNI3* or *CACNA1C* on day 0. Relative expression of *TNNI1* was not significantly different between LD1 (0.02-fold) and LD2 (0.025-fold), however both had significantly lower relative expression compared to PB. A similar trend was observed for day 0 *RYR2* expression, with LD1 and LD2 exhibiting significantly lower expression (0.6- and 0.67-fold, respectively) relative to PB. *CASQ2* was the only gene on day 0 to be significantly differentially expressed between LD1 and LD2 (0.85- and 1.63-fold, respectively), with LD2 having significantly greater *CASQ2* expression than both LD1 and PB.

#### Day 14

Relative *TNN1* expression was significantly lower on day 14 in both LD1 (0.05-fold) and LD2 (0.01-fold) relative to PB. *TNNI3* expression was significantly lower in both LD1 (0.73-fold) and LD2 (0.11-fold) relative to PB, with LD2 having significantly lower relative expression than LD1 as well. PB’s higher relative expression of these myofibrillar genes indicate that PB hiPSC-CMs are more cardiogenic than LD1 and LD2 hiPSC-CMs on a transcription level, with LD2 hiPSC-CMs being the least cardiogenic. Furthermore, the *TNNI3*:*TNNI1* ratio is a metric of myofibrillar maturity, with a lower *TNNI3*:*TNNI1* ratio associated with less mature CMs^22,23^. Normalizing day 14 *TNNI3*:*TNNI1* expression to day 0 for each population, PB (3.28×10^-4^) had a significantly higher *TNNI3:TNNI1* compared to LD1 (1.31×10^-4^) and LD2 (8.77×10^-5^), further demonstrating PB hiPSC-CMs’ greater maturity relative to LD1 and LD2 hiPSC-CMs (**Fig. S11**). Relative expression of *RYR2* on day 14 was significantly lower in LD2 (0.32-fold) compared to both PB and LD1 (1.13-fold), which were not significantly different from one another. Relative *CASQ2* expression was significantly greater in LD1 (1.92-fold) compared to both PB and LD2 (0.75-fold). *CACNA1C* expression was significantly lower in LD2 (0.29-fold) relative to both PB and LD1 (0.82-fold), which were not significantly different from each other. Notably, while there are significant differences in relative cardiac gene expression on days 0 and 14, these relative differences were not conserved from day 0 through day 14, indicating that baseline levels of gene expression in an hiPSC lineage is not determinant of upregulation of those same genes after differentiation.

These results indicate that lineage identity impacts cardiac differentiation outcomes, particularly in areas of CM yield and gene expression. Furthermore, ClonMapper can be used to identify non-cardiogenic lineages, and lineages maintain their differentiation potential when differentiated in isolation.

### TagSeq Highlights Differential Expression

Using analysis of TagSeq sequenced RNA from PB, LD1, and LD2 on days 0, 5, and 14 of differentiation, clustermaps were generated based on genes that are significantly upregulated (**Fig. 5A**) or downregulated (**Fig 5B**). Examining upregulated genes, *RPS4Y1* is consistently upregulated in PB and downregulated in LD1 and LD2 at all timepoints (**Fig. 5A(i)**). Though *RPS4Y1* is not known to be involved in CM differentiation or function^24^, these results identify *RPS4Y1* as a potential underlying driver of the poor differentiation outcomes of LD1 and LD2 compared to PB. Genes related to CM myofibrillar structure (*MYL3, MYH7B, MYOM1*)^25–27^ and function (*PPP1R14C, ASB2*)^28–31^ are most upregulated in day 14 PB and LD1, but far less expressed in LD2 (**Fig. 5A(ii)**), highlighting them as particularly critical markers of cardiac differentiation and development, even more so than currently established. While not preferentially expressed by CMs, genes in **Fig. 5A(iii**) have reported involvement in influencing CM maturation and CVD^32–37^, and are upregulated in all day 14 samples but are most upregulated in LD2 and least in LD1, indicating they may contribute to non-cardiogenic differentiation outcomes.

**Figure 5:**
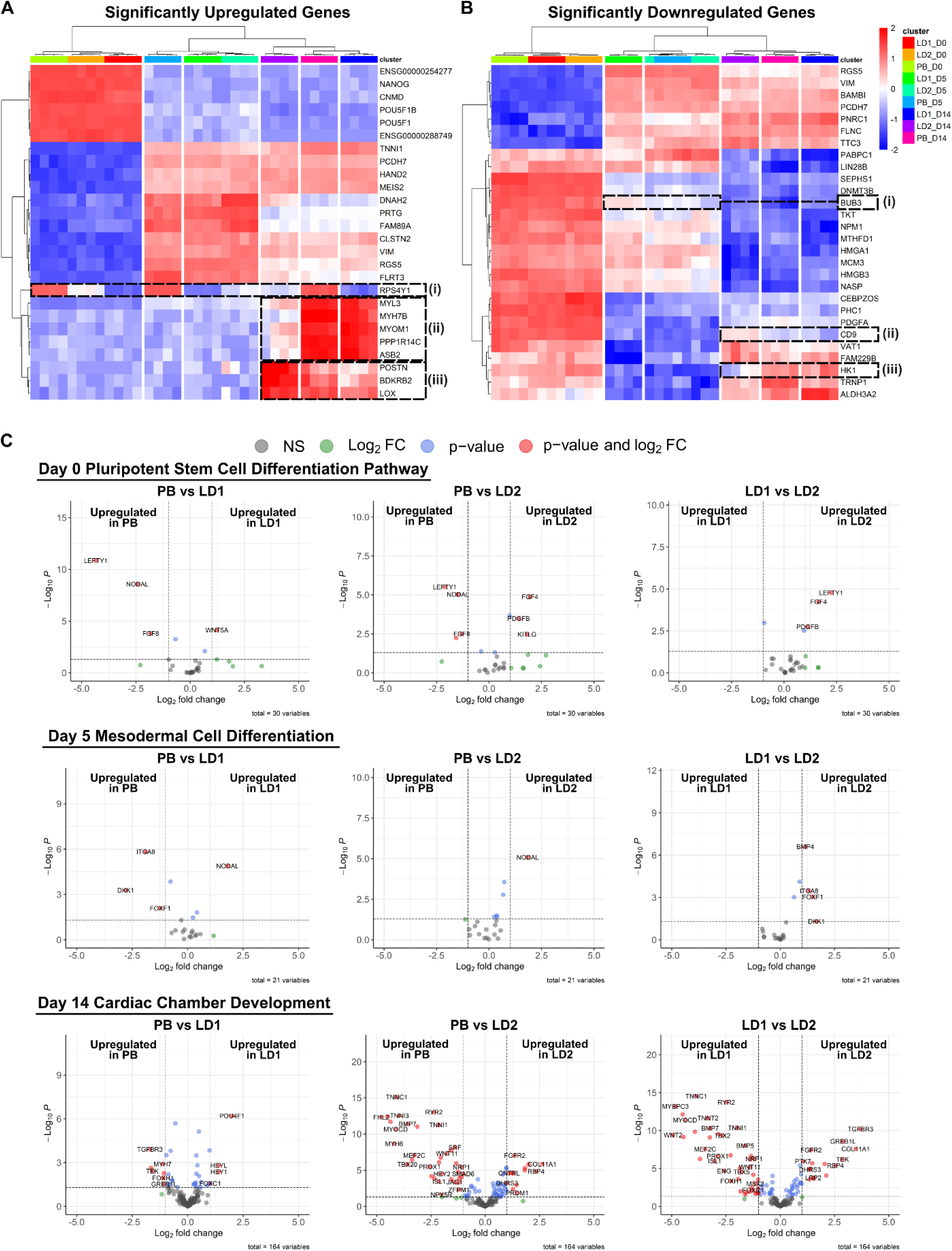
Analysis of Tagseq data from PB, LD1, and LD2 RNA isolated on days 0, 5, and 14 of differentiation. A clustermap of significantly upregulated genes (**A**) and significantly downregulated genes (**B**) shows PB, LD1, and LD2 biological replicates cluster together, indicating differences in gene expression between the populations. Furthermore, key genes that are more selectively up- or downregulated in specific populations (**A(i-iii**), **B(i-iii)**) highlight potentially underlying drivers and markers of poor cardiac differentiation outcomes. Volcano plots showing genes in relevant gene sets that are significantly differentially expressed in pairwise comparisons between PB, LD1, and LD2 on days 0, 5, and 14 (**C**) show differential expression of key genes.

Examining downregulated gene clustering (**Fig. 5B**), day 5 LD1 and LD2 biological replicates inter-cluster, indicating potential overlap in downregulation of genes on day 5 that contributes to poorer cardiac differentiation outcomes. *BUB3* was only downregulated in LD1 and LD2 on day 5 (**Fig. 5B(i)**) and was recently reported as a potential regulator of extensive *OBSCN* alternative splicing during cardiac muscle development^38^. Lower *BUB3* expression may have disrupted regulation of *OBSCN* splicing during differentiation of LD1 and LD2, resulting in poorer outcomes. *CD9* was downregulated in day 14 PB and LD1 but upregulated in LD2 (**Fig. 5B(ii)**), with *CD9* having been reported to worsen pathological cardiac hypertrophy in mice^39^. This prior evidence of *CD9*’s involvement in the heart along with our results mark *CD9* as a potential indicator of poor differentiation outcomes. *HK1* was downregulated in certain day 14 LD2 replicates while upregulated in all PB and LD1 replicates (**Fig. 5B(iii)**). Given that hexokinases have reported involvement in cardioprotection^40^, our results indicate that downregulation of *HK1* may be a marker of poor differentiation outcomes. Furthermore, generating additional clustermaps of key pathways that regulate hiPSC cardiac differentiation (**Fig. S12**) revealed that only the P53 signaling pathway resulted in discrete clustering of biological replicates, suggesting that genes regulated by P53 signaling result in more distinct transcriptomic differences between PB, LD1, and LD2.

Lastly, specific relevant gene sets on days 0 (PSC differentiation pathways), 5 (mesodermal cell differentiation), and 14 (cardiac chamber development) were used to analyze gene expression and generate volcano plots based on log2(FC) and statistical significance (p-value) of the gene foldchange (**Fig. 5C**). Examining day 0 plots, *NODAL*, *FGF8*, and *LEFTY1* were significantly upregulated in PB relative to LD1 and LD2. *NODAL* signaling is vital for initiating in vitro cardiac differentiation^41,42^, and *FGF8* is critical in the formation of the primitive streak^43^, suggesting that lower expression of these two genes in non-cardiogenic lineages LD1 and LD2 on day 0 may contribute to their poorer outcomes. Though *LEFTY1* inhibits the Nodal signaling pathway in the developing heart^44^, its upregulation in PB suggest that *LEFTY1* upregulation may not be as inhibitory of Nodal signaling during in vitro directed differentiation. Examining day 5 pairwise comparisons, *NODAL* is significantly upregulated in LD1 and LD2 compared to PB. Given that prolonged *NODAL* signaling in cardiac mesodermal cells inhibits subsequent CM specification while inhibition of *NODAL* after cardiac mesoderm formation improves CM yield in human PSC cardiac differentiations^42,43^, these results confirm LD1 and LD2 are less cardiogenic, with upregulated *NODAL* being a potential cause. Furthermore, since inhibition of *BMP4* at this stage of differentiation similarly improves CM yields^42^, upregulation of *BMP4* in LD2 relative to LD1 corroborates LD2 being the least cardiogenic lineage. For day 14 pairwise comparisons, upregulation of *MYH7* in PB relative to LD1 corroborates clustermap results (**Fig. 5A**), and upregulation of *TNNI3, TNNI1*, and *RYR2* in PB and LD1 relative to LD2 corroborate our qPCR results and the conclusion that LD2 is the least cardiogenic lineage.

## DISCUSSION

Herein we adapted ClonMapper cell barcoding to investigate hiPSC-CM differentiation. We demonstrated that hiPSCs labeled with ClonMapper barcodes remain pluripotent and can differentiate into hiPSC-CMs with no negative impact on CM yield or ECC metrics, with the retention of a functional GCaMP6f transgene demonstrating that ClonMapper does not interfere with pre-existing transgenes, meaning it could be used in conjunction with reporter hiPSC lines. Furthermore, while ClonMapper barcodes were silenced during differentiation, they were still retained in nuclei and could be used to assess clonal lineage dynamics, as Jiang et al. 2022 demonstrated clonal dynamics can be successfully evaluated through hiPSC cardiac differentiation even with transcriptionally silenced barcodes^45^. Tracking lineages through differentiation revealed that enrichment of PB lineages in CM or non-CM populations was impacted by lineage identity, suggesting intrinsic differences between hiPSC lineages resulted in lineage-specific outcomes. This was in contrast to prior findings on hiPSC-CM cell barcoding from Jiang et al. 2022^45^, though it is important to note that Jiang and colleagues used a different cell barcoding platform, hiPSC line, and cardiac differentiation protocol. Tracking lineages through differentiation also revealed that lineages enriched in the CD140a+ population on day 5 of differentiation did not consistently differentiate into hiPSC-CMs by day 14 despite CD140a being a reported marker of cardiogenic differentiation trajectory^20^, highlighting the value of using cell barcoding to identify useful markers to help improve differentiation methods.

Using ClonMapper’s recall function we isolated two clonal lineages that were enriched in non-CMs, LD1 and LD2, to establish two homogeneous, single-clone hiPSC populations. Our inability to isolate cardiogenic lineages warrants further investigation in future studies adapting ClonMapper or other functional genetic barcodes to hiPSC differentiation workflows. qPCR revealed both LD1 and LD2 had significantly different levels of cardiac gene expression compared to PB hiPSCs. Furthermore, differentiating PB, LD1, and LD2 revealed significant differences between the three populations in hiPSC-CM yield and gene expression levels, indicating that distinct hiPSC lineages respond differently to the same cardiac differentiation stimuli. TagSeq analysis of PB, LD1, and LD2 throughout differentiation further demonstrated that distinct hiPSC lineages from the same hiPSC line behave differently, as the three populations had significant transcriptomic differences throughout differentiation. Additionally, using ClonMapper to live-isolate the non-cardiogenic LD1 and LD2 helped identify *RPS4Y1* as a potential underlying driver of poor cardiac differentiation outcomes and highlighted several other genes as pathways as potential markers and drivers of poor differentiation outcomes that warrant further investigation.

These findings demonstrate that ClonMapper can be used to successfully identify and isolate hiPSC lineages with low cardiac differentiation efficiency, and that different hiPSC lineages respond differently to the same cardiac differentiation protocol. Lastly, our inability to isolate lineages enriched in hiPSC-CMs highlights the need for additional optimization when applying ClonMapper workflows to hiPSC.

The data presented here support further investigation of specific genes and pathways as drivers of heterogeneous differentiation outcomes and use of hiPSC barcoding in cardiac and other differentiation pathways. Furthermore, examining clonal lineages in different hiPSC lines could elucidate whether clonal lineage-specific responses to differentiation cues originate from reprogrammed somatic cell type, emerge during reprogramming process, or arise after hiPSC reprogramming, particularly given prior reports of lineage-dependent iPSC reprogramming outcomes^46,47^. Our findings highlight the need to investigate what drives success in cardiac differentiation of hiPSC lineages, how these differences might inform experimental design, and whether they are experimentally tunable.

## CONCLUSION

We have demonstrated that ClonMapper can be used to establish a barcoded hiPSC population without negatively impacting cardiac differentiation outcomes, track clonal lineages through differentiation, identify lineage-specific outcomes, and isolate lineages with specific responses to differentiation cues. Single-lineage hiPSC populations can be established and differentiated, with these homogenous populations displaying cardiac differentiation outcomes that are distinct from the heterogeneous barcoded population, and serving to identify specific genes and pathways as potential markers or drivers of poor cardiac differentiation outcomes which warrant future investigation. Different cardiac differentiation approaches have different focuses on how to improve cardiac differentiation outcomes such as medium formulations, substrate composition and architecture, bioreactor design, or defined coculture with non-CMs^6,48^. Exploring these approaches is critical to developing hiPSC-CMs into a clinically viable resource. However, accounting for lineage-specific behavior is a potential missing piece that could inform the use of all the previously discussed methods, with the results from cell barcoding with ClonMapper supporting this hypothesis. Utilizing cell barcodes will enable the tracking of lineage-specific responses to various differentiation stimuli for individual hiPSC lineages, which will elucidate underlying mechanisms that drive changes in differentiation outcomes and help improve hiPSC cardiac differentiation yield and functionality, and ultimately improve the clinical scalability and relevance of hiPSC-CMs.

## METHODS

### hiPSC Culture

WTC11 (GM25256, Coriell Institute) hiPSCs modified to express GCaMP6f (a gift from Dr. Bruce Conklin) were the primary hiPSCs used in experiments, and from which PB, LD1, and LD2 populations were generated. hiPSCs were cultured in Essential 8 (E8) medium (Thermo Fisher, A1517001) with daily medium exchanges on vitronectin (Thermo Fisher, A14700) coated tissue culture plates. hiPSCs were passaged when ∼85% confluent (every 3-4 days).

### Cardiac Differentiation

hiPSC-CMs were differentiated according to methods adapted from Lian et al 2013^49^ discussed in supplementary materials (**Methods S1**).

### Immunostaining for Microscopy and Cytometry/FACS

The full list of antibodies and respective dilutions is available in supplementary materials (**Table S1**) along with immunostaining methods used (**Methods S4**).

### qPCR

Total RNA was isolated from cells then reverse transcribed into cDNA used for qPCR. Relative mRNA expression was quantified using the ΔΔC_T_ method with TATA-binding protein (*TBP*) as the endogenous control to calculate ΔC_T_ for all genes. All primers used are listed in the supplementary materials (**Table S2**), along with more detailed methodologies (**Methods S5**).

### ClonMapper Barcode, Plasmid, and Lentivirus Production

Barcodes, plasmids, and lentivirus were prepared as previously reported^50,51^ and discussed in supplementary materials (**Methods S2**).

### ClonMapper Barcode Integration

#### Lentiviral Titer

Viral titers were performed for each ClonMapper lentivirus batch to determine the amount of virus needed to achieve a multiplicity of infection (MOI) ≤ 0.1, according to published ClonMapper methods^51^ discussed in supplementary materials (**Methods S3**).

#### Barcode Transduction

hiPSCs were transduced and prepared for FACS similar to **Methods S3**, with the exception that the volume of virus solution added to hiPSCs was the predetermined volume needed to achieve an MOI ≤ 0.1.

### Labeling hiPSCs with ClonMapper Barcodes

hiPSCs were transduced with 27.9 µL/well of virus to achieve an MOI ≤ 0.1, based on a lentiviral titer of 1.33×10^6^ Titering Units/mL (**Fig. S1**) and cell count of 412,500 cells/well on the day of transduction. Barcoded hiPSCs were sorted using “Single Cell” purity on a Sony MA900 into vitronectin-coated 96-well plates containing E8 with 10 µM Y-27632 (Selleck Chemicals, S1049) and 1:100 antibiotic-antimycotic (Sigma Aldrich, A5955-100ML). Sorted, barcoded hiPSCs were expanded and passaged until populations could seed a full 6-well plate. Sorted wells were not mixed during expansion, so each 6-well plate represented a distinct heterogeneously barcoded population. Barcoded hiPSCs were cryopreserved at 1M-2M cells/mL in 9:1 fetal bovine serum:dimethyl sulfoxide (Sigma Aldrich, F0926 and 472301). hiPSCs thawed from liquid nitrogen underwent three passages prior to use in experiments.

#### hiPSC ClonMapper Recall

ClonMapper recall was done according to published ClonMapper methods^51^, as discussed in supplementary materials (**Methods S6**), with recalled hiPSCs sorted as was described for barcode-transduced hiPSCs.

#### Barcoded hiPSC-CM Functionality

ECC metrics of hiPSC-CMs were quantified using time-stacks of GCaMP6f fluorescence images captured at 33 fps from day 14 control and barcoded hiPSC-CMs to generate fluorescent signal traces. These traces were analyzed using a MATLAB pipeline for quantifying duration, synchronicity, and decay constant of calcium signals in hiPSC-CMs we have previously reported on (**Fig. S3**)^15^. Changes in relative expression of cardiac genes were analyzed via qPCR performed with cDNA generated on days 0, 7, and 14 of differentiation, with non-barcoded controls used as the reference control group at each timepoint to quantify ΔΔC_T_ values.

### Establishing a Parental Barcoded Population and Assessing Clonal Dynamics

Genomic DNA (gDNA) was extracted from heterogeneously barcoded populations using a PureLink Genomic DNA Mini Kit (Invitrogen, K182001), underwent PCR amplification and purity assessments according to published ClonMapper methods^50^, then was sequenced on an Illumina NextSeq 500 (Genomic Sequencing and Analysis Facility (GSAF) at UT Austin, Center for Biomedical Research Support. RRID: SCR_021713) with 5M reads per sample. Barcode reads were extracted from FASTQ files using Pycashier^52^, a computational package for analyzing ClonMapper barcode data, to quantify the relative abundance of each unique barcode and the Shannon Diversity of each population in R. gDNA was extracted from the selected Parental barcode (PB) population on days 0, 4, 7, 10, and 20 of differentiation, with gDNA then amplified and sequenced, with barcode reads extracted using Pycashier then analyzed in R.

### Identifying Lineage Differentiation Potential

gDNA was extracted from differentiating PB hiPSCs on days 0, 5, and 14 of differentiation. Cells were sorted on a Sony MA900 on days 5 and 14 after staining for CD140a/PDGFRα (cardiac mesoderm marker^20^) and SIRPα (hiPSC-CM marker^16^), respectively, with gDNA extracted immediately after sorting, with ∼550K-1.2M cells sorted per subpopulation (CD140a+, CD140a-, SIRPα+, SIRPα-). Extracted gDNA was amplified, purified, then sequenced on an Illumina NovaSeq as pooled libraries with 5M reads per sample. Barcode reads were extracted from FASTQ files with Pycashier and analyzed in R. Barcode sequences not found in all experimental replicates were filtered out, and then relative abundance of each lineage was calculated and normalized to log2(CPM) values.

Relative change in abundance of each lineage was determined by calculating the log2(FC) of each barcode sequence in day 5 CD140a+, day 5 CD140a-, day 14 SIRPα+, and day 14 SIRPα- populations relative to day 0 undifferentiated hiPSCs, with log2(FC) values then averaged across experimental replicates.

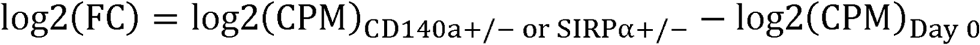

### ClonMapper Recall of PB Lineages

Custom recall plasmids cardiogenic and non-cardiogenic PB lineages were made according to published ClonMapper methods^50^ discussed in supplementary materials (**Methods S7**). ClonMapper recall was performed as previously described with recalled cells expanded and cryopreserved, and gDNA extracted, amplified, purified, then Sanger sequenced (Eton Bioscience, ABI 3730xl DNA Sequencers) to assess purity of the recalled and expanded populations.

Isolated populations confirmed as pure, single-lineage populations were differentiated into hiPSC-CMs, with RNA isolated on days 0, 5, and 14 of differentiation for qPCR and Tagseq. For qPCR, PB was used as the reference sample for determining ΔΔC_T_ values for all samples on days 0 and 14. Time-stack images of beating hiPSC-CMs’ GCaMP6f fluorescence were captured for analyzing ECC and cells were immunostained to assess CM yield on day 14.

### Tagseq of PB, LD1, and LD2

Tagseq libraries were sequenced using a NovaSeq 6000 SR100 at the UT Austin GSAF and preprocessed by the Bioinformatics Consulting Group at the University of Texas at Austin. Reads were then processed using the nf-core RNAseq pipeline^53,54^. Briefly, reads were aligned to human reference genome GRCh38 using ‘STAR’, before quantifying gene counts using ‘Salmon’^55,56^.

Differentially expressed genes were determined using edgeR’s quasi-likelihood method^57^. Differentially expressed genes were defined to have an adjusted p-value <0.05 (Benjamini-Hotchberg) and log-fold-change of ±1. Genes, ranked by log-fold-change, were used for gene-set enrichment analysis with fgsea package^58^ in R and the Gene Ontology Biological Processes set, WikiPathways, Kegg, Reactome, Pathway Interactions Database, and MSigDB Hallmark gene sets^59–65^. Significantly enriched pathways were defined as having adjusted p-values <0.05. Gene counts and associated code can be found at https://github.com/ZoldanLab/CM_barcoding.

### Statistical Analyses

Bar graphs represent data using mean values with standard deviation error bars. CM yield and ECC were analyzed with student’s t-tests for non-barcoded vs barcoded comparisons and with one-way ANOVA for PB vs LD1 vs LD2. qPCR was analyzed with two-way ANOVA. TagSeq statistics are as previously described and documented in associated code.

## RESOURCE AVAILABILITY

### Lead Contact

Additional information on methods, resources, and reagents should be directed to the lead contact, Dr. Janet Zoldan.

### Materials Availability

All materials in this study are available from commercial vendors listed in the methods or from the authors.

### Data and Code Availability

MATLAB code is available upon request from the lead contact. Data, code, and reagents lists for ClonMapper are available at https://docs.brocklab.com/clonmapper/. Data and code for analysis of barcode gDNA next-generation sequencing and TagSeq are available at https://github.com/ZoldanLab/CM_barcoding. All authors adhered to FAIR and CARE data management practices.

## Acknowledgements

We thank Michael Cotner for assistance with plasmid preparation, and Dr. Nikhith Kalkunte, Dr. Brett Stern, and Anna McClain for assistance with hiPSC cell culture. We acknowledge and appreciate support from funding sources: National Heart, Lung, and Blood Institute F31 Grant (5F31HL170717-03; S.S.).

## Declarations of Interest

The authors declare no competing interests.

## Notes

### Competing Interest Statement

The authors have declared no competing interest.

