## Supplemental Materials for "Cell Barcoding Reveals Lineage-dependent Outcomes in hiPSC Cardiac Differentiation"

**SUPPLEMENTARY METHODS**

*Methods S1: hiPSC Cardiac Differentiation*. 3 days before differentiation was to be initiated (day -3), hiPSCs were seeded on Geltrex-coated (Thermo Fisher, A1413202) tissue culture plates at 35,000 cells/cm^2^ in E8 medium with E8 replaced every 24 h. After 3 days (day 0), differentiation was initiated by exchanging medium with RPMI 1640 (Thermo Fisher, 11875135) with B27 Supplement minus insulin (Thermo Fisher, A1895601; RB-) supplemented with 12 µM CHIR99021 (LC Labs, C-6556). After differentiation was initiated, subsequent medium exchanges until day 9 were performed within 1 hour of when initiation occurred. On day 1, medium was replaced with RB-. On day 3, a 1/2 medium exchange was performed, replacing spent medium with RB- supplemented with 10 µM IWP-2 (Tocris Bioscience, 3533) such that final concentration of IWP-2 in each well would be 5 µM. On day 5, medium was replaced with RB-. On day 7, medium was replaced with RPMI 1640 medium with B27 supplement (Thermo Fisher, 17504044; RB+). On day 8, medium was replaced with RB+ without glucose (RPMI 1640 without glutamine, Thermo Fisher, 21870076). On day 9, medium was replaced with RB+, with subsequent medium exchanges occurring with RB+ every 48-72 h. Spontaneously beating hiPSC-CMs were observable on day 8 but were often most apparent on day 10 due to the 24 h glucose deprivation on day 8.

*Methods S2: ClonMapper Barcode, plasmid, and lentivirus production.* Briefly, two oligonucleotides (IDT) were generated (CROPseq-PrimeF-BgL-BsmBI: GAGCCTCGTCTCCCACCGNNNNNNNNNNNNNNNNNNNNGTTTTGAGACGCATGCTGCA

and CROPseq-RevExt-BgL-BsmBI: TGCAGCATGCGTCTCAAAAC) and used in a 4X extension reaction to generate the barcode library insert. This insert was then ligated into a pre-digested backbone, Crop-Seq-BFP-WPRE-TS-hU6-BsmBI (Addgene, 137993), in a 50X golden gate reaction with BsmBI (NEB, R0739S). Following golden gate assembly, plasmid was electroporated into E. coli in 500 mL of 2xYT medium containing 100 µg/mL carbenicillin and incubated overnight at 37ºC. Bacterial cells were pelleted by centrifugation at 6,000g at 4ºC for 15 min and plasmid DNA was extracted using a QIAGEN Plasmid Plus Midi kit (QIAGEN, 12943). ClonMapper lentivirus was generated as previously described ^28,29^. Briefly, 48 h prior to transfection 2.5 x 10^5^ HEK293T cells were seeded in each well of a 6-well plate. Each well was transfected for 16 h using Lipofectamine 3000 (ThermoFisher, L3000001) with 1.5 µg psPAX2 (Addgene, 12260), 0.4 µg VSV-G (Addgene, 14888), and 5 µg ClonMapper barcode library plasmid. 48 h following transfection viral containing supernatant was concentrated ~20X using 30,000 MWCO PES concentrators (Sartorius, VS2022).

*Methods S3: Determining ClonMapper Viral Titer.* E8 was prepared with 10 µg/mL polybrene and increasing amounts of ClonMapper lentiviral solution then added to hiPSCs 24 h after they were passaged at 22,000 cells/cm^2^, with medium replaced with fresh E8 after 15-18 h. After 24 h the cells were detached, rinsed in DPBS twice, resuspended in sorting buffer (DPBS with 0.5% FBS and 2 mM EDTA), then strained through 40 µm mesh into 5 mL sorting tubes. Flow cytometry was performed on a Sony MA900 Multi-Application Cell Sorter (Center for Biomedical Research Support Microscopy and Imaging Facility at UT Austin. RRID:SCR_021756) to measure the abundance of blue fluorescent protein (BFP) signal, as successful ClonMapper barcode transduction results in BFP production. The viral titer volumes and their corresponding percent BFP^+^ signal were plotted to determine the relationship between the amount of virus and MOI (**Fig. S1**) which was used to calculate the amount of virus needed to achieve the target MOI ≤ 0.1 in subsequent transductions.

*Methods S4: Immunostaining*. For imaging adherent cells, were fixed in 4% paraformaldehyde, which was then neutralized with 300 mM glycine in DPBS. Fixed cells were permeabilized with 0.2% Triton X-100, blocked with blocking buffer (0.1% Tween-20 and 1% bovine serum albumin (BSA)), and then incubated in primary antibody solutions overnight at 4°C. Cells were rinsed twice with blocking buffer then incubated in secondary antibody solutions for 1 hr at room temperature in the dark, then rinsed and stained with DAPI. Full list of antibodies and respective dilutions available in supplementary information (**Table S1**).

For live-staining of surface proteins, cells were detached then resuspended in surface blocking buffer (DPBS with 1% BSA) and kept on ice for 10 min, then pelleted and resuspended in 1% BSA with 1/10 of the suspension aliquoted as a negative control. Diluted primary antibody solution was added to non-negative control cells and kept on ice on a shaker for 1 h. Cells were then pelleted and washed twice in 1% BSA. Diluted secondary antibody was then added to all samples and incubated on ice for 30 min. Cells were then pelleted and resuspended/washed in sorting buffer twice before being strained through 40 µm mesh caps into 5 mL sorting tubes and kept protected from light.

For intracellular protein staining, cell pellets were resuspended in 4% PFA and incubated on ice for 10 min before being rinsed in DPBS with glycine and pelleted, twice. Fixed cells were pelleted then permeabilized in DPBS with 0.1% Triton X-100 on ice for 10 min, after which the cell suspension was washed in DPBS then pelleted once more. Cells were then resuspended in blocking buffer and kept on ice on a shaker for 30 min. The remaining steps were then identical to methods for staining surface proteins described above.

*Immunostaining to Assess Pluripotency*. Barcoded hiPSCs were immunostained for pluripotent markers using Oct4 and SOX2 primary antibodies and goat-anti-rabbit-AF488 secondary antibody. Barcoded hiPSCs differentiated into hiPSC-CMs were immunostained using an anti-cardiac troponin T (cTnT) primary antibody and donkey anti-mouse-AF488 secondary antibody.

*Immunostaining to Assess hiPSC-CM Yield.* Both barcoded and non-barcoded control hiPSC-CMs underwent cardiac differentiation. Differentiated cells were then immunostained for flow cytometry with either anti-cTnT or anti-SIRPα/CD147A primary antibodies, then goat-anti-mouse-AF647 secondary antibody.

*Methods S5: RNA Isolation, Reverse Transcription, and RT-qPCR.* Total RNA was isolated from cells using a RNeasy Mini Kit (Qiagen, 74104) and reverse transcription was performed using a High-Capacity cDNA Reverse Transcription Kit (Applied Biosystems, 4368814) on a Verti 96-well Thermocycler (Applied Biosystems) with the following thermal profile: 10 min at 25°C, 120 min at 37°C, and 5 min at 85°C.

Quantitative PCR was performed using PrimeTime qPCR Primers (Integrated DNA Technologies) and PowerUp SYBR Green (Thermo Fisher) on a StepOnePlus Real-Time PCR System (Applied Biosystems). The reactions were activated for 2 min at 50°C then 2 min at 95°C, and then underwent forty amplification cycles: 15 s at 95°C and 60 s at 60°C.

*Methods S6: ClonMapper Recall in hiPSCs.* Recall control (RC) hiPSCs were seeded on vitronectin-coated wells at 22,000 cells/cm^2^ and transfected 24 h later by dropwise addition of medium comprising E8 with Opti-Mem (Gibco, 31985062) and Lipofectamine 3000 (Invitrogen, L3000001) supplemented with plasmid solutions. A control well for fluorescent gating was left un-transfected, a control well for normalizing transfection efficiency was transfected with a green fluorescent protein (GFP) expression plasmid, and a third well was transfected with the two plasmids needed to induce recall activity, with total amount of plasmid DNA kept the same between the transfected wells. The two plasmids used to induce ClonMapper recall were a dCas9-VPR plasmid and a recall plasmid containing a sequence complimentary to the sequence in the RC hiPSCs that enables dCas9-VPR to bind the plasmid such that downstream transcription of a GFP gene occurs. Medium was exchanged 15-18 h after transfection with E8, and the next day transfected cells were lifted using Accutase and prepared for sorting as previously described. hiPSCs expressing GFP due to recall plasmid transfection were sorted using “Single Cell” purity of a Sony MA900 into a 5 mL sorting tube containing sorting buffer.

*Methods S7: ClonMapper Recall of hiPSCs from PB Population.* Reverse complements of the barcode sequence in each target lineage were used to generate three pairs of overlapping oligos (Thermo Fisher, 10336022) that were annealed into DNA blocks then ligated to form a 3X-barcode array which was then ligated into a recall plasmid backbone using a Golden Gate assembly reaction. Competent *E. coli* were transformed with the Golden Gate product and plated on carbenicillin-containing agar plates. Plasmid DNA was extracted from transformed bacteria (QIAGEN, 12123) then sequenced (Plasmidsaurus, Whole Plasmid Sequencing) to confirm barcode insertion, with correctly assembled plasmids undergoing an additional expansion via culture of *E. coli* transformed with the sequenced plasmid.

**SUPPLEMENTARY FIGURES**

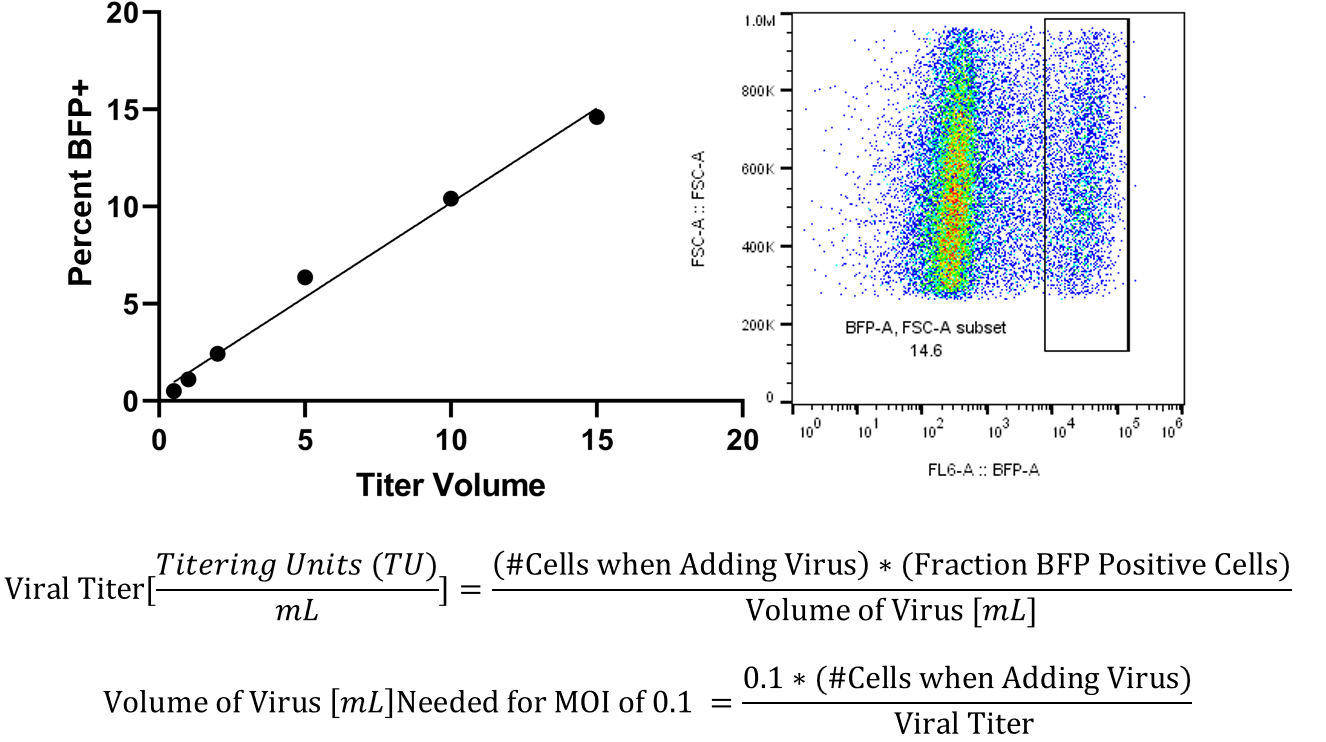

%BFP+ = 0.9632*[volume of virus] + 0.4316

**Supplementary Figure S1:** Viral titer assay was performed to determine volume of viral solution needed to achieve an MOI ≤ 0.1. The percentage of BFP⁺ cells was plotted against the volume of viral solution added (left). BFP expression was quantified by flow cytometry, with representative flow cytometry data shown for the sample transduced with 15 µL of virus solution (right). Viral titer was calculated using the above equations with cell counts from a sacrificial replicate well.

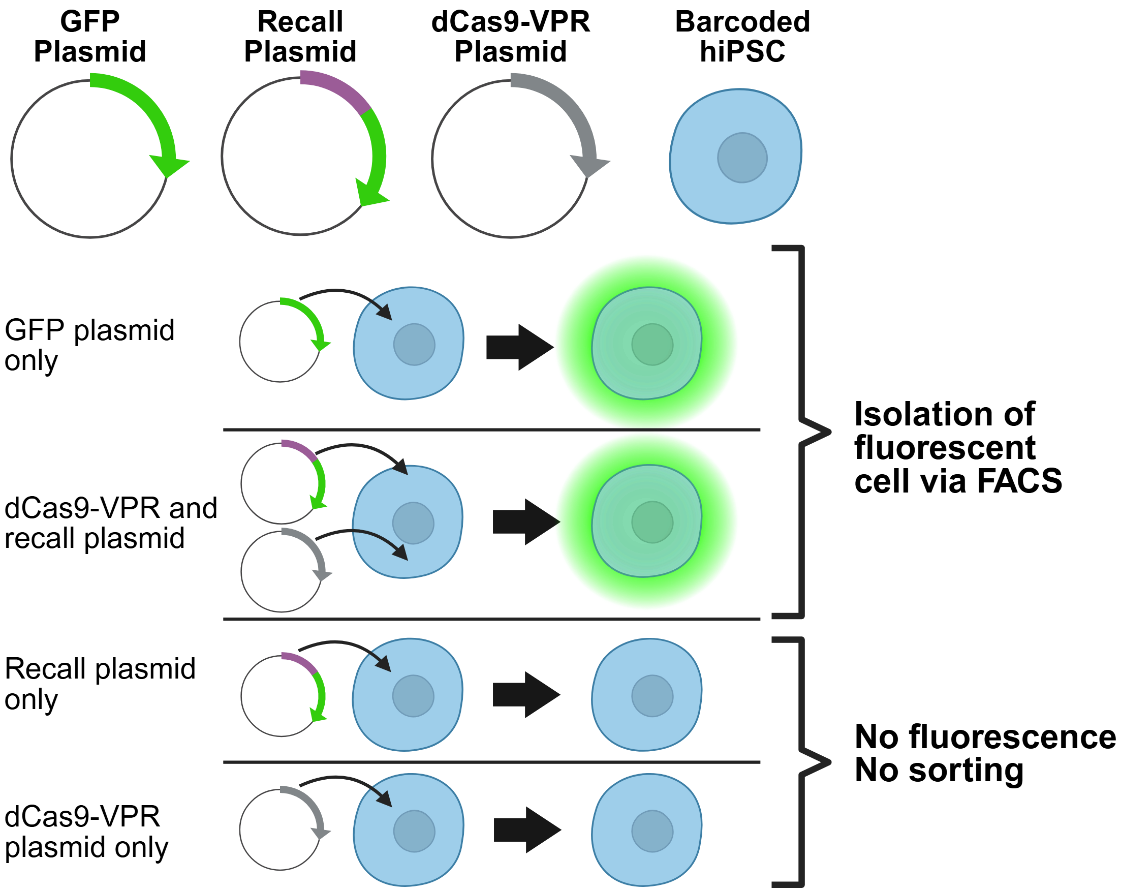

**Supplementary Figure S2.** Diagram depicting the transfections necessary for green fluorescent protein (GFP) expression to occur in barcoded cells. ClonMapper’s lineage recall takes advantage of gRNA’s role in moderating Cas protein activity by transfecting barcoded cells with a dCas9-VPR plasmid and a “recall plasmid” that encodes a ClonMapper barcode sequence complement and a GFP gene. If the sequence on the recall plasmid is complementary to the barcode sequence of a transfected cell, then dCas9-VPR binds to the recall plasmid and activates transcription of GFP. Fluorescence in barcoded hiPSCs was actuated by a constitutive GFP expression plasmid or by the interaction of the dCas9-VPR plasmid, recall plasmid, and barcode within the barcoded hiPSC.

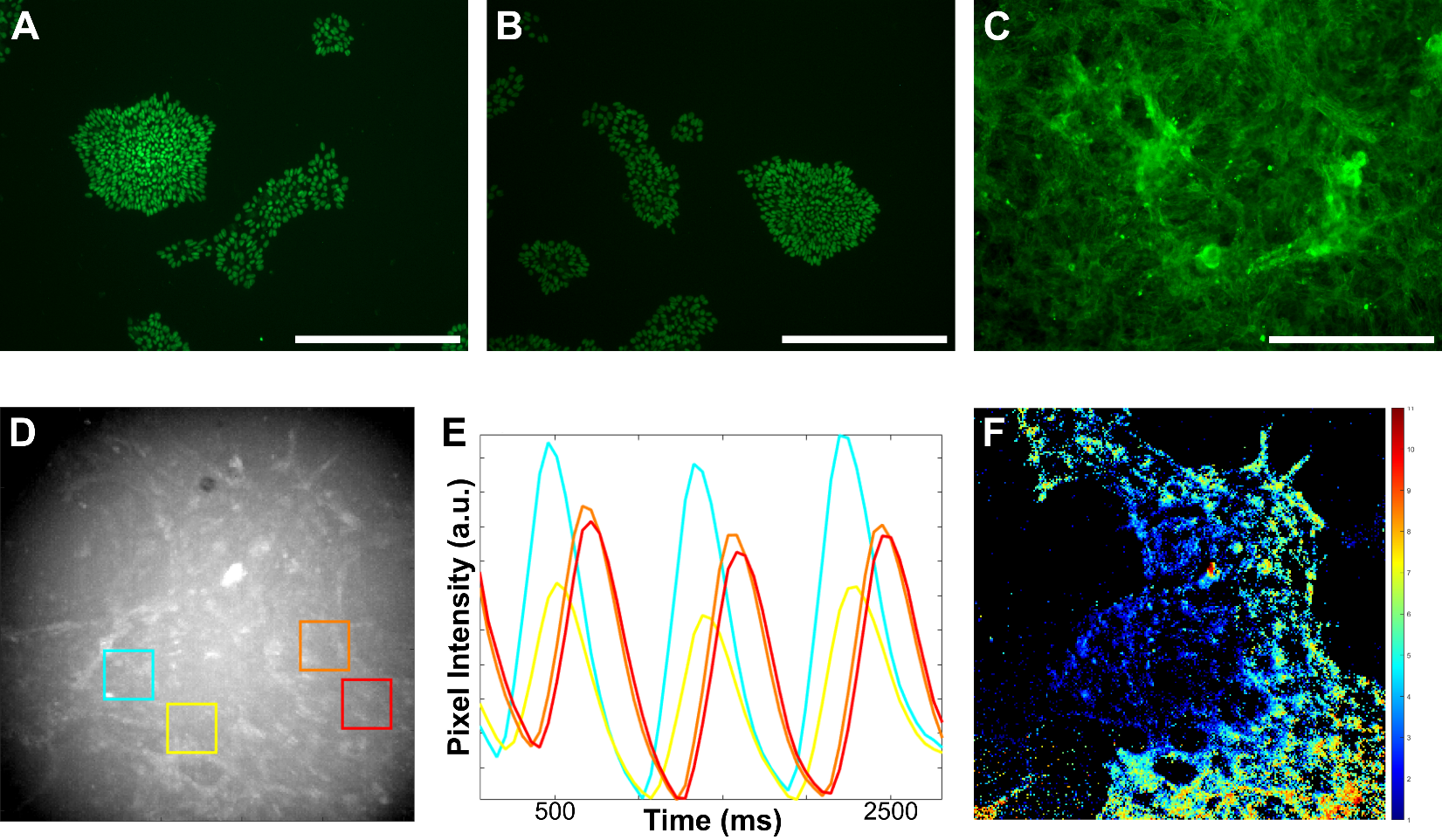

**Supplementary Figure S3:** hiPSCs labeled with ClonMapper barcodes remain pluripotent and able to successfully differentiate into hiPSC-CMs while retaining prior genetic modifications. Representative fluorescent images showing that barcoded hiPSCs express canonical pluripotency markers SOX2 (**A**) and Oct4 (**B**) (BFP signal overlap not shown due to fixation and permeabilization causing loss of BFP produced after transduction). Representative fluorescent images demonstrating that barcoded hiPSCs successfully differentiated into cTnT expressing cardiomyocytes (**C**). The use of hiPSCs that express GCaMP6f enables calcium excitation-actuated GFP fluorescence in differentiated hiPSC-CMs, a property retained after barcoding. This allows excitation-contraction coupling to be visualized via fluorescence and quantified using an in-house MATLAB analysis pipeline (**D**). Time-stacks of images of beating cells were analyzed to extract metrics such as beating traces (**E**) and synchronicity (**F**). All scale bars are 400 µm.

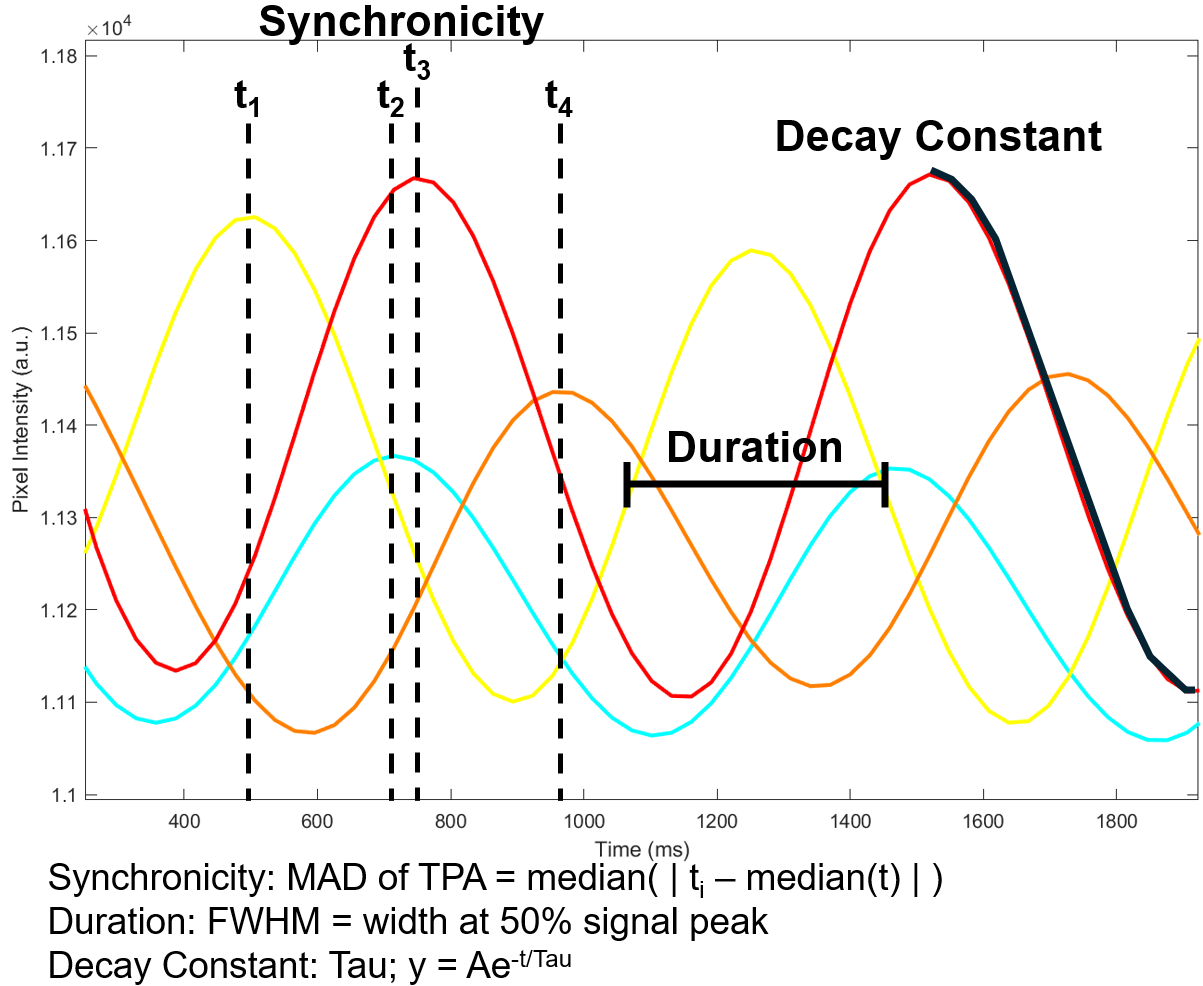

**Supplementary Figure S4:** Representative signal traces depicting the excitation-contraction coupling metrics computed by the Zoldan lab’s MATLAB analysis pipeline, generated from time-stack images capturing calcium-actuated GCaMP6f fluorescence from spontaneously beating hiPSC-CMs. MAD of TPA: median-absolute-deviation of time-of-peak-arrival; FWHM: full-width at half-max.

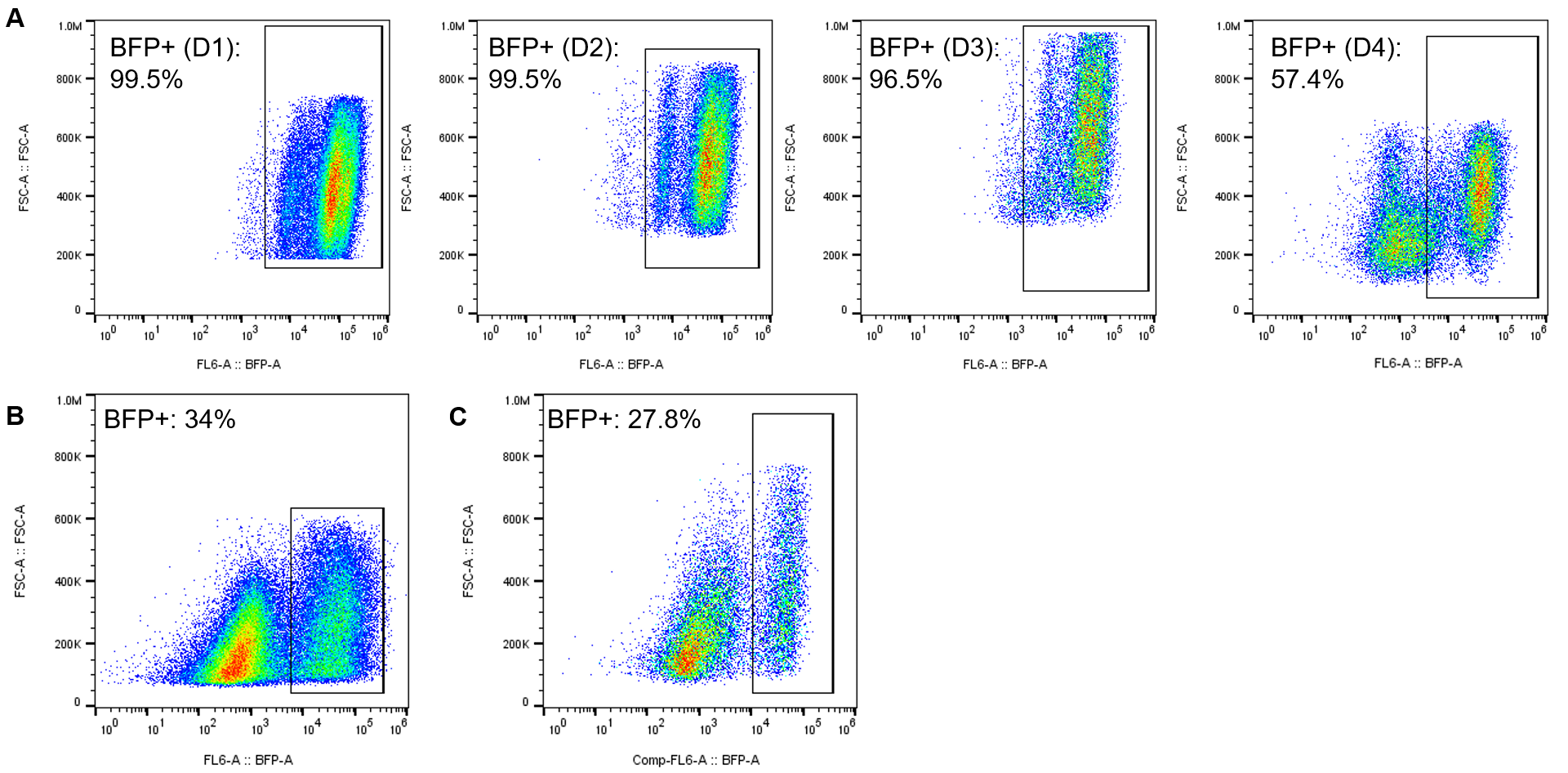

**Supplementary Figure S5:** Flow cytometry data showing reduction in fraction of BFP-expressing cells after Wnt inhibition on day 3 of cardiac differentiation in GCaMP6f-expressing WTC11 hiPSCs (**A**). Similar silencing was observed in cardiac differentiation of unmodified (no GCaMP6f expression) WTC11 hiPSCs (**B**) and CD34^+^ endothelial progenitor differentiation of DF19-9-11T.H hiPSCs (**c**). Endothelial progenitor differentiation was adapted from Jalilian & Raimes 2020^19^.

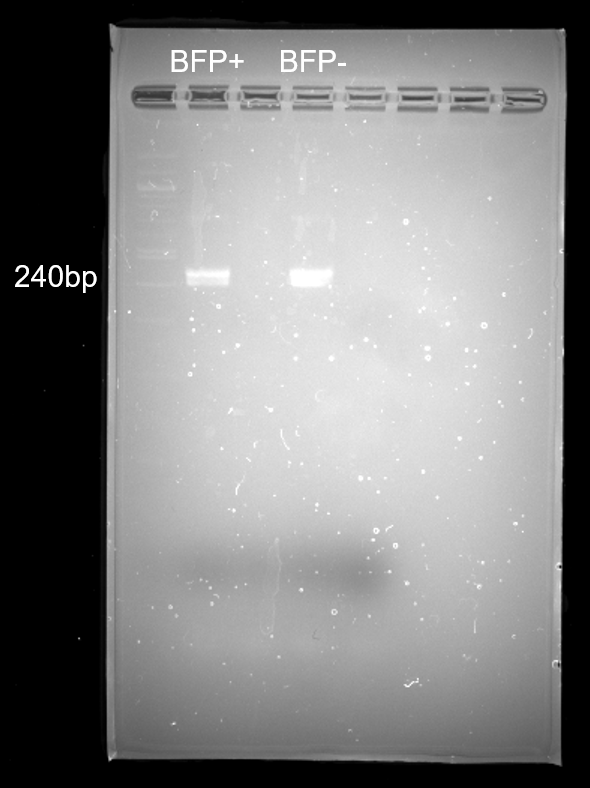

**Supplementary Figure S6:** Agarose gel showing electrophoresed genomic DNA from BFP+ and BFP- cells differentiated from an entirely BFP+ hiPSC population both retain the target barcode sequence that is ~240 base pairs (bp) long. Accordingly, ClonMapper barcodes are retained in all differentiated cells even if BFP is silenced and no longer transcribed, resulting in a loss of BFP signal.

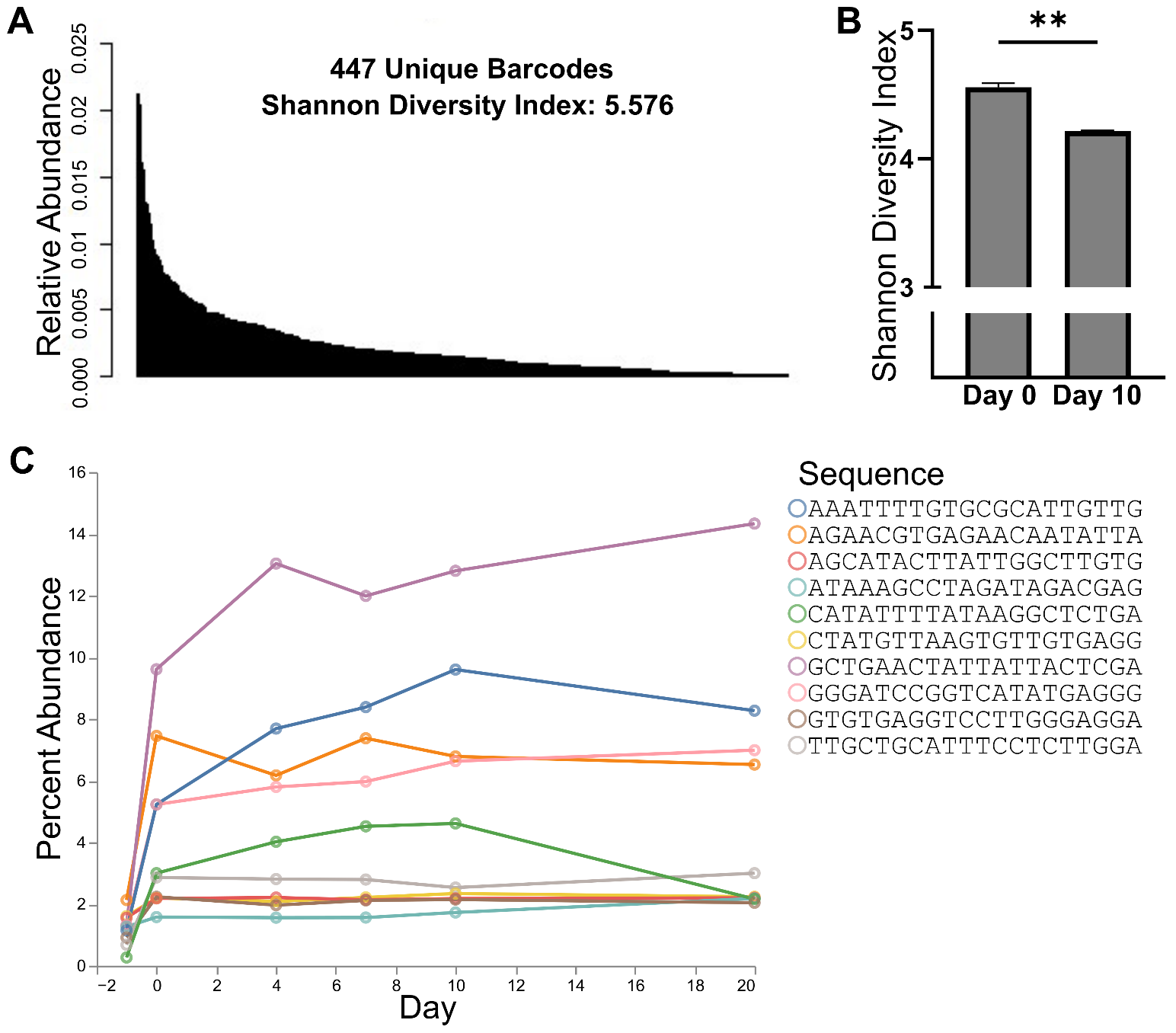

**Supplementary Figure S7:** Validation that ClonMapper parental barcoded (PB) hiPSC population is diverse enough to capture clonal dynamics through cardiac differentiation and that significant changes in barcode population diversity and relative abundance occur over time. PB hiPSCs cryopreserved for further experimentation have a high diversity with a broad distribution without any lineages dominating the population (**A**). Shannon diversity of different replicates shown to decrease through differentiation, with a statistically significant difference in diversity between thawed PB hiPSCs prior to differentiation (Day 0) and differentiated hiPSC-CMs (Day 10) (**B**; n=3, ** indicates *p* ≤ 0.01). Snapshot of one experimental replicate showing changes in relative abundance as percent frequency of the 10 most abundant lineages in the undifferentiated population (**C**).

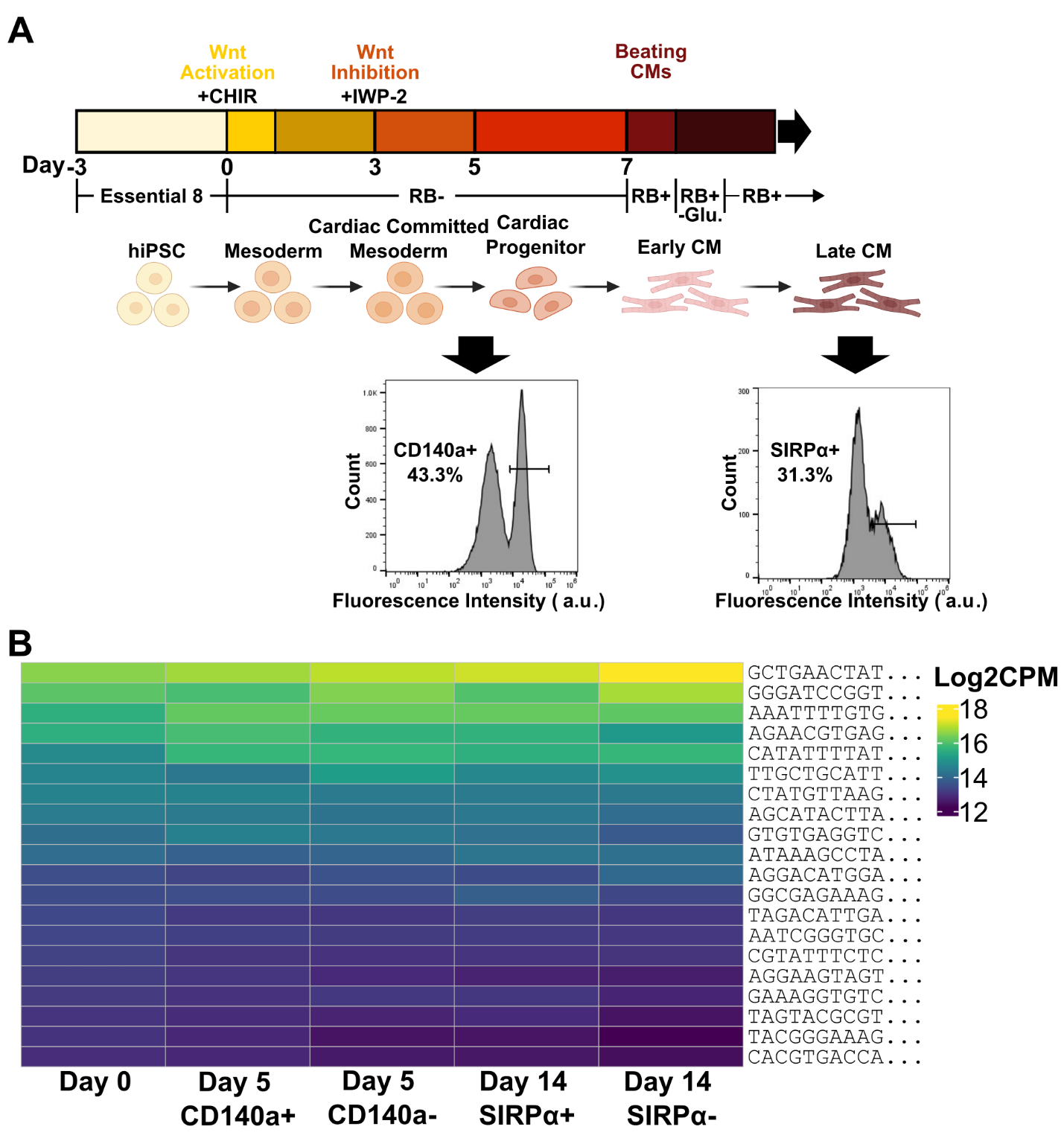

**Supplementary Figure S8:** Cardiac differentiation protocol timeline. PB hiPSCs were differentiated and sorted on days 5 and 14 of differentiation after immunofluorescent staining for CD140a and SIRPα, which are cardiac mesoderm and hiPSC-CM markers, respectively (**A**). Heatmap of the 20 most abundant day 0 PB lineages based on log2(CPM) of abundance in the day 5 and day 14 sorted populations (CD140a+/- and SIRPα+/-, respectively) (**B**).

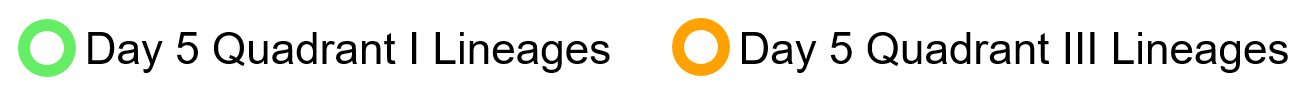

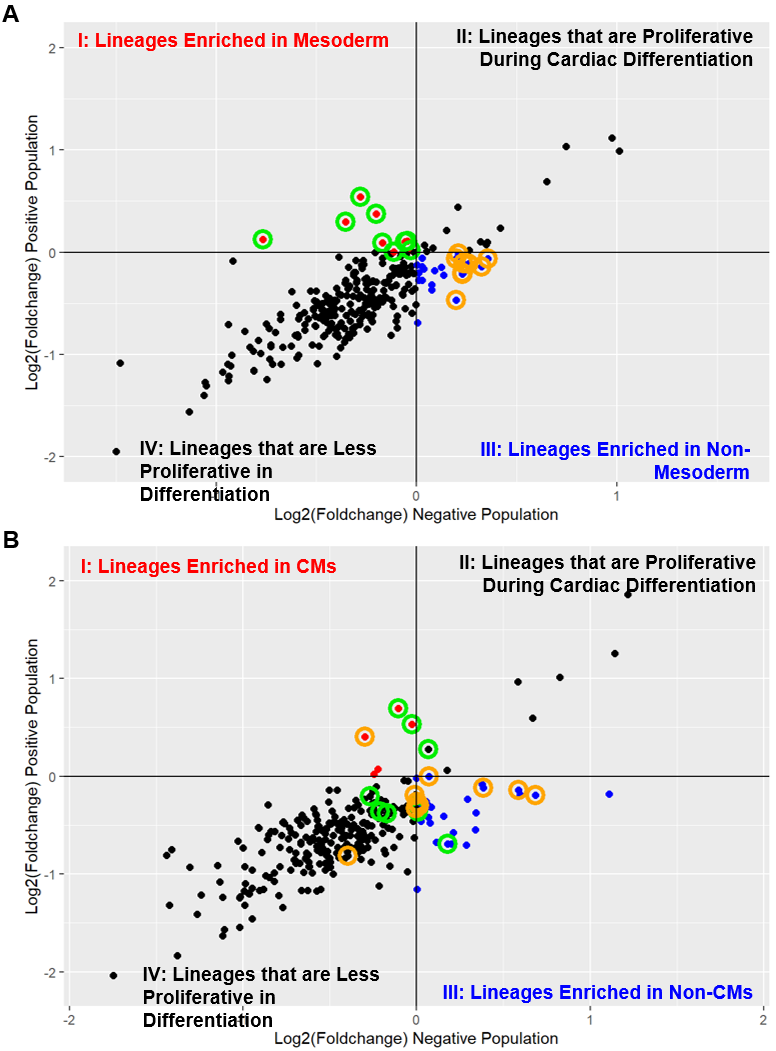

**Supplementary Figure S9:** Log2(foldchange) of lineage abundance in positive and negative sorted populations plotted against each other for day 5 (**A**; n = 6) and day 14 (**B**; n = 6), with log2(foldchange) being relative to abundance of lineages on day 0. Tracking lineages enriched in mesodermal, CD140a+ cells (points circled in green) and lineages enriched in non-mesodermal, CD140a- cells (points circled in orange) from day 5 in the day 14 SIRPα sorted samples shows no relationship between CD140a and outcome of cardiac differentiation.

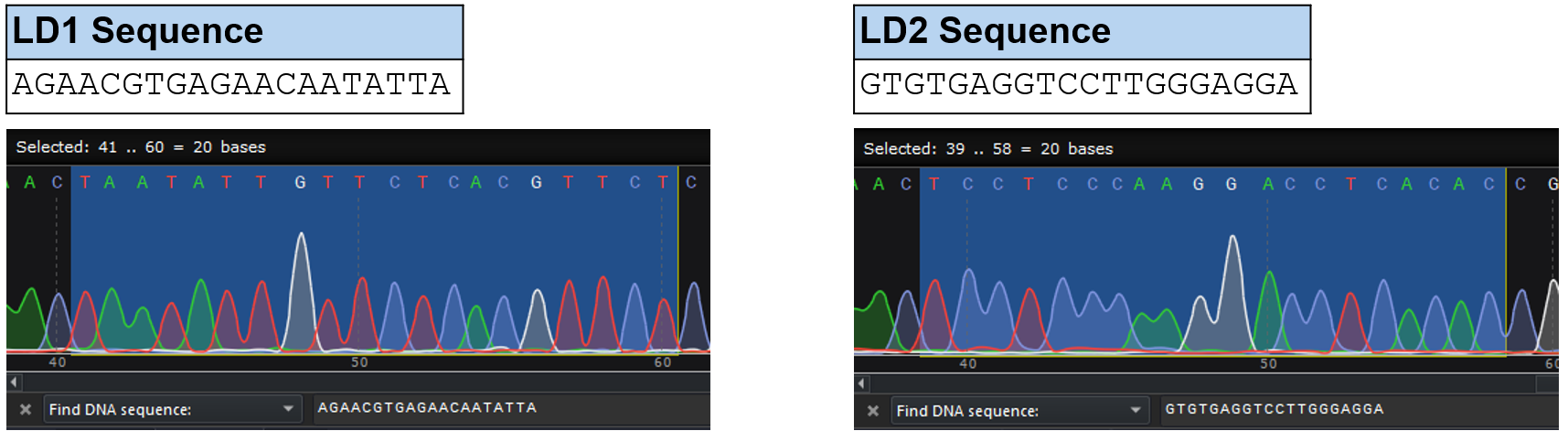

**Supplementary Figure S10:** Sanger sequencing traces demonstrating that DNA replicated from genomic DNA extracted from LD1 and LD2 cells sorted using ClonMapper recall contain reverse complements of the target lineage barcode sequence. Accordingly, these isolated populations are comprised solely of the target lineage.

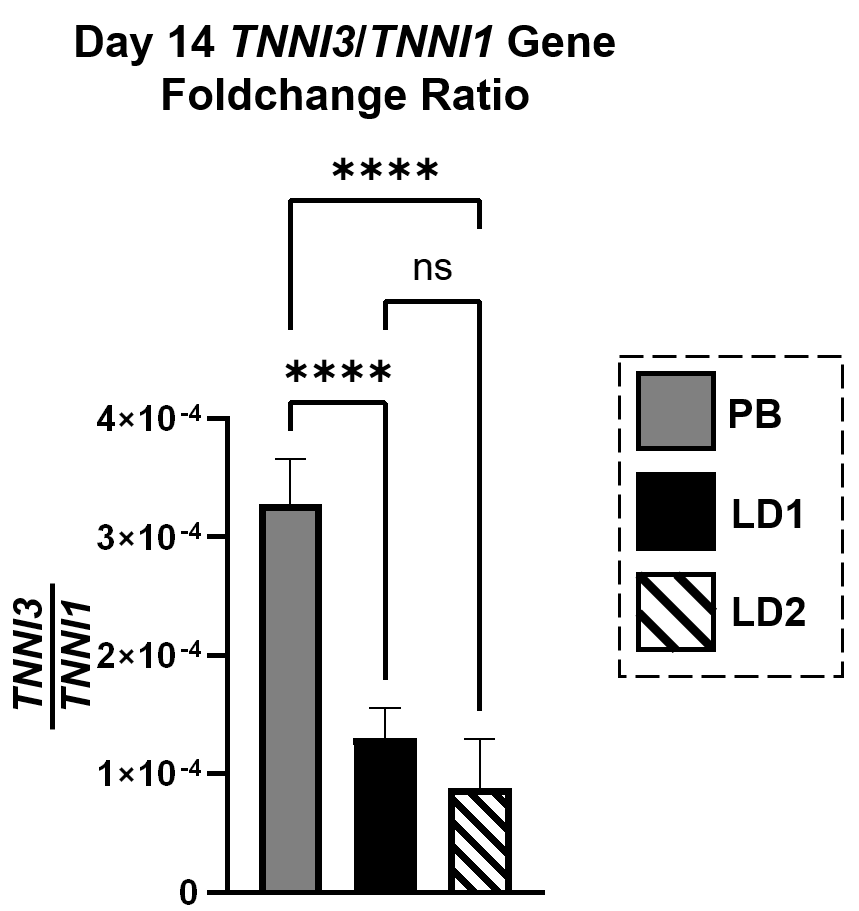

**Supplementary Figure S11:** Ratio of TNNI3 to TNNI1 gene foldchange in day 14 differentiated cells relative to day 0 undifferentiated hiPSCs.TNNI3:TNNI1 ratio is an indicator of cardiomyocyte maturity, with a higher ratio being associated with greater CM maturity.

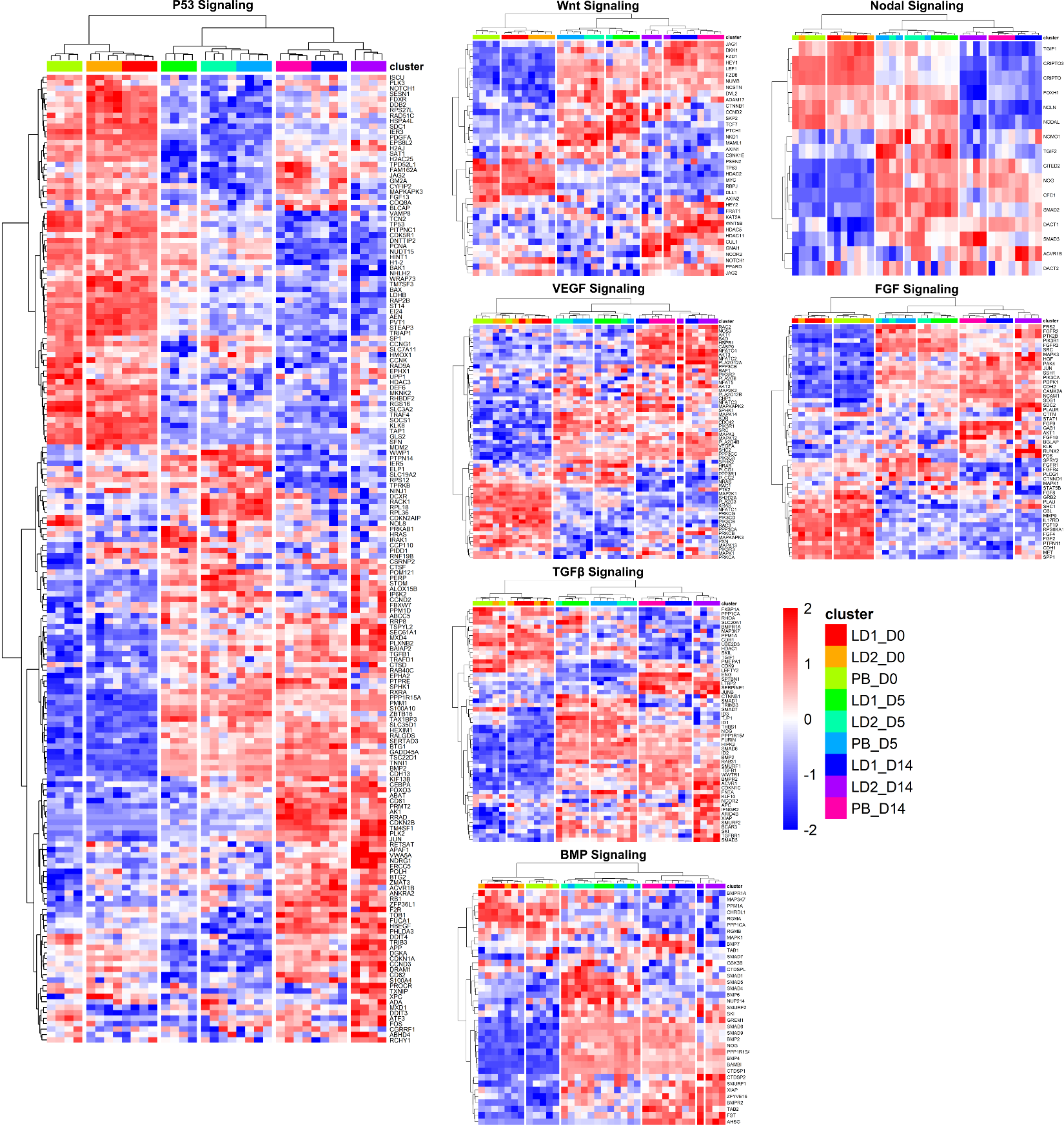

**Supplementary Figure S12:** Clustermaps of key pathways involved in cardiomyocyte differentiation. The interspersion of PB, LD1, and LD2 biological replicates resulting in lack of discrete clustering of distinct populations through all timepoints suggests that the signaling regulated by these pathways may not be the underlying cause of differences between PB, LD1, and LD2 populations in cardiac differentiation.

**SUPPLEMENTARY TABLES**

**Supplementary Table S1: Immunostaining Antibodies and Dilutions**

|  | **Antibody** | **Source** | **Dilution** |
| --- | --- | --- | --- |
| **Primary Antibodies** | Mouse-anti-cardiac troponin T | Abcam, ab8295 | Microscopy: 1:1000  Flow Cytometry/FACS: 1:300 |
|  | Mouse-anti-SIRPα | BioLegend, 323802 | Flow Cytometry/FACS: 1:50 |
|  | Rabbit-anti-Oct4 | Applied StemCell, ASA-0110 | Microscopy: 1:6 |
|  | Rabbit-anti-SOX2 | Applied StemCell, ASA-0120 | Microscopy: 1:6 |
| **Secondary Antibodies** | Goat-anti-rabbit-AF488 | Abcam, ab150077 | Microscopy: 1:500 |
|  | Goat-anti-mouse-AF647 | Jackson ImmunoResearch, 115-605-003 | Flow Cytometry/FACS: 1:400 |
|  | Donkey-anti-mouse-AF488 | Abcam, ab150105 | Microscopy: 1:200 |
| **Fluorophore-Conjugated Antibodies** | Human-anti-CD140a-PE | Miltenyi, 130-131-952 | Flow Cytometry/FACS: 1:100 |

**Supplementary Table S2: qPCR Primer Sequences**

| **Gene Name** | **Assay ID** | **Primer Sequence** |
| --- | --- | --- |
| TBP | Hs.PT.58v.39858774 | 5'-GCTGTTTAACTTCGCTTCCG-3' |
|  |  | 5'-CAGCAACTTCCTCAATTCCTG-3' |
| TNNI1 | Hs.PT.58.1261401 | 5'-GAACAAGGTGCTGTCTCACT-3' |
|  |  | 5'-CCAGCATTCCTTGGCCTT-3' |
| TNNI3 | Hs.PT.58.23287465 | 5'-CAGGACTTGTGCCGACAG-3' |
|  |  | 5'-CGCTTAAACTTGCCTCGAAG-3' |
| RYR2 | Hs.PT.58.502763 | 5'-CAGAGTTCGCACAGTAACAGT-3' |
|  |  | 5'-CAGCCAATCTCCAGTCCATT-3' |
| CACNA1C | Hs.PT.58.14979004 | 5'-CTTCGAGTACCTGATGTTCGTC-3' |
|  |  | 5'-GTTGAGGATGTTCATGGCGAT-3' |
| CASQ2 | Hs.PT.56a.219158 | 5'-CAATACTGACAACCCCGATCTG-3' |
|  |  | 5'-TTCACCACCCCAATCTGTG-3' |
